# Targeting SUFU-mediated GLI activity alleviates ciliopathic phenotypes

**DOI:** 10.64898/2026.10.23.760487

**Authors:** Ching-Fang Chang, Yanfen Yang, Kari M. Brown, Daniel T. Rasicci, Timothy T. Nguyen, Kevin A. Peterson, Samantha A. Brugmann

**Affiliations:** Division of Developmental Biology, Department of Pediatrics, Cincinnati Children’s Hospital Medical Center, Cincinnati, Ohio 45229, United States; University of Cincinnati, College of Medicine, Department of Pediatrics, Cincinnati, Ohio, 45229 United States; Division of Plastic Surgery, Department of Surgery, Cincinnati Children’s Hospital Medical Center, Cincinnati, Ohio 45229, United States; The Jackson Laboratory, Bar Harbor, Maine 04609, United States

## Abstract

Ciliopathies are incurable, pleiotropic diseases that result from defective cilia. Several ciliopathic phenotypes arise from improper processing and activity of GLI transcription factors, yet attempts to rescue these phenotypes by overexpressing activator or repressor GLI isoforms have proven unsuccessful. Here, we demonstrate that modulating SUFU-GLI interactions restores GLI transcriptional activity and rescues ciliopathic phenotypes *in vivo*. As previous work identified increased accumulation of SUFU in ciliopathic conditions, we hypothesized that exploiting SUFU’s epistatic relationship to the cilium could provide a novel therapeutic strategy for ciliopathies. Genetic introduction of either a single null allele (*Sufu^+/-^*) or point mutation allele (*Sufu^D159R^*), which modulates SUFU-GLI binding, into a ciliopathic mouse model revealed that both interventions improved midfacial phenotypes; however, *Sufu^D159R^* provided a more complete rescue by significantly improving mandibular phenotypes as well. Molecular examination of this genetic modification revealed restored GLI3 binding to DNA, GLI3 interaction with transcriptional cofactors, and GLI3-directed transcription of target genes, collectively driving the rescue of numerous ciliopathic phenotypes. These findings reveal cilium-independent functions for SUFU and establish that modulating SUFU-GLI interactions bypasses the need for ciliary GLI processing. Together, these data provide a promising therapeutic approach for ameliorating ciliopathic pathologies.

## INTRODUCTION

Ciliopathies comprise a large group of syndromes that affect the human nervous, cardiovascular, respiratory, gastrointestinal, nephrogenic, visual, and skeletal systems (1). Ciliopathies are due to disruptions in the form or function of the primary cilium, a microtubule-based extension on the surface of most cells which is essential for molecular, mechanical, osmotic, and chemical signal transduction. The diversity of phenotypes associated with ciliopathies has hampered the development of treatment options for this disease class.

The primary cilium functions as a signaling hub, most notably playing an essential role in transducing the Hedgehog (HH) pathway (2). HH receptors, Patched1 (PTCH1) and Smoothened (SMO), localize to the cilium (3–5). When HH signaling is active, HH ligands bind to PTCH1 on the surface of the cilium, this binding alleviates PTCH1-mediated repression of SMO, and activated SMO then works through the microtubule motor protein Kinesin (6, 7) and inhibition of Protein Kinase A, by mechanisms that remain unclear, resulting in processing of full-length GLI (GLIFL) into a functional GLI activator (GLIA) (8, 9). When HH signaling is inactive, the full-length isoforms of GLI2 and GLI3 are processed into a truncated GLI repressor (GLIR) (10–12). Full-length GLI1, by contrast, is not processed into a truncated repressor form because it lacks both a processing determinant domain and a repressor domain, and thus only functions as an obligate activator of the pathway. When the cilium is absent or non-functional, all GLIFL forms are unable to function as activators and GLIR formation is largely reduced (5, 13, 14). Consequently, tissues reliant upon GLIA present with a loss of HH phenotype and those reliant on GLIR present with a gain of HH phenotype when the cilium is compromised (15, 16).

We previously reported that in a conditional ciliopathic mouse model (*Kif3a^fl/fl^;Wnt1Cre*), facial development was compromised due to a gain of HH activity in the frontonasal prominence (FNP) and loss of HH activity in the mandibular prominences (MNPs) (15, 16). The gain-of-function phenotype in the FNP was partially rescued by expressing GLI3R, and the loss-of-function phenotype in the MNP was partially rescued by expressing GLI2A. This partial rescue, however, was concomitant with exacerbation of the phenotype in the other facial prominence (i.e., improvement of the FNP correlated with exacerbation of the MNP phenotype, and vice versa). Thus, we began to examine alternative approaches to modulate GLI activity as potential therapeutic options for ciliopathies.

We and others have reported increased expression of Suppressor of fused (SUFU) and accumulation of SUFU-GLI protein complexes in mouse models of ciliopathies (13, 15). SUFU is regarded as a negative regulator of the HH pathway due to the close phenotypic resemblance of *Sufu^-/-^* and *Ptch1^-/-^* embryos (17). SUFU acts as a binding partner and modulator of all three GLI transcription factors (TFs), regulating their sequestration in the cytoplasm, ciliary trafficking, nuclear transport, and transcriptional activity (17–22). SUFU interacts physically with GLI proteins through several residues and motifs. First, Sufu-Interacting N-terminal (SIN) domain in all three GLI proteins is comprised of the RSSL (23) and SYGHLS motifs (24). The SIN domain binds to motifs in the C-terminal region of SUFU and to the D159 residue (25). The interaction between the SIN domain and SUFU has been ascribed multiple functions including GLI stabilization, cytoplasmic retention, ciliary-localization, and a target site for phosphorylation by multiple protein kinases (23, 25–28). Second, the SUFU interacting C-terminal (SIC) domain (27) in GLI proteins binds to the N-terminal region of SUFU, including but not limited to the D159 residue. The interaction between the SIC domain and SUFU is required for transcriptional activity (27). Third, the SUFU-binding repressor domain (SR) is only present in GLI2 and GLI3 and binds to both the folded SIC domain and to the residues in the N-terminus of SUFU (22, 23, 29, 30).

Several mechanisms for SUFU-mediated GLI repression have been proposed (13, 14, 20, 21). Initially, SUFU was thought to act as a cytoplasmic constraint for the nuclear translocation of GLI TFs (13, 19, 31), thus preventing transcriptional activation of GLI targets. However, evidence indicated that SUFU can translocate to the nucleus with GLIR and recruit the SAP18-SIN3A corepressor complex (21) which is required for transcriptional repression. Similarly, it was reported that SUFU must dissociate from the full-length transcriptional activator (GLIA) to function (22). More recent studies, however, suggest that upon HH activation, SUFU-GLIFL complexes are recruited from the cytoplasm to the primary cilium and accumulate at its distal tip, where GLIFL is processed into GLIA. At the ciliary tip, HH stimulation promotes dissociation of the SUFU-GLI complex, releasing GLI to enter the nucleus and activate transcription (5, 31). The ciliary tip thus serves as a key site for the regulation of SUFU-GLI complexes. However, whether full dissociation of SUFU from GLI is required for GLI to activate transcription has remained unresolved. SUFU has also been reported to remain associated with GLIA and translocate with it into the nucleus to activate GLI target genes (22). Thus, despite being an essential modulator of GLI activator and repressor activity, the mechanisms of SUFU activity and interactions with GLI TFs remain unclear.

Considering the increased expression of SUFU and accumulation of SUFU-GLI protein complexes in ciliopathic mouse models, along with the fact that SUFU is epistatic to the cilium (14, 32) and contributes to GLIA and GLIR transcriptional activity, we hypothesized that SUFU might be a viable target for treating ciliopathies. Herein, we examine a role for SUFU in restoring GLI3 activity under ciliopathic conditions. Our data indicate that mutating a single amino acid in SUFU (*Sufu^D159R^*) (29, 30) restored GLI3 binding to DNA, binding to cofactors, subsequent expression of GLI target genes, and rescued ciliopathic phenotypes. These findings suggest the cilium is required for GLI binding to DNA, as well as interactions with transcriptional cofactors, and that modulating SUFU-GLI interactions can bypass the need for ciliary processing. Together, these data provide a promising target for future therapeutic approaches to ameliorate ciliopathic pathologies.

## RESULTS

### SUFU interacts with full-length GLI3 to form transcriptional complexes

SUFU interacts with GLIs through several residues and motifs (22, 23, 29, 30). Previous characterization of the ciliopathic mouse model *Kif3a^fl/fl^;Wnt1Cre* revealed excessive amounts of SUFU and SUFU-GLI complexes (15). To investigate the biological significance of this interaction within a craniofacial context in greater depth, we first examined the interaction of SUFU with full-length GLI3 (GLI3FL), the major GLI transcription factor in craniofacial development, using AlphaFold3 (33). In the predicted murine SUFU-GLI3 complex structure, the previously reported SIN, SIC, and SR domains (23), as well as the five zinc-finger DNA-binding domains of GLI3 were present. Additionally, the characteristic N-and C-terminal lobes, as well as the D159 residue of SUFU were present (Fig. 1A-A’). Modeling predicted SUFU was in contact with the SIN, SIC and SR domains of GLI3 and surrounded by large, disordered regions (Supplementary Fig. 1A). A chord diagram of inter-chain contacts showed that SUFU engaged GLI3 predominantly through these three domains, which together accounted for 178 of the 185 high-confidence residue-residue contacts (SIN, 100; SIC, 51; SR, 27; contact probability ≥ 0.5; Fig. 1A and Table S1). Notably, the model placed D159, the only functionally important residue of the SUFU acidic patch, in direct, high-confidence contact with the SIN region, specifically the SYGHLS motif (Ser333-His336; contact probability 0.88-0.92; Table S1), which was the most heavily engaged interface, consistent with prior structural and functional studies (30). These SUFU-GLI3 contact regions were also conserved in GLI1 and GLI2 (Supplementary Fig. 1B). To test this interaction *in vivo*, we performed co-immunoprecipitation in E11.5 MNPs from mice expressing endogenously and N-terminally FLAG-tagged GLI3 (*Gli3^BioFLAG^* allele) with anti-FLAG antibody. Although GLI3R was more prevalent than GLI3FL in the MNP (Fig. 1B, input), SUFU co-immunoprecipitated with more GLI3FL than GLI3R (Fig. 1B, IP). Thus, although SUFU is regarded predominantly as a repressor of the HH pathway that recruits corepressor complexes, these data supported the suggestion that SUFU also associated with the GLI3FL activator and may play a role in GLI3-mediated gene activation (22).

**Figure 1.**
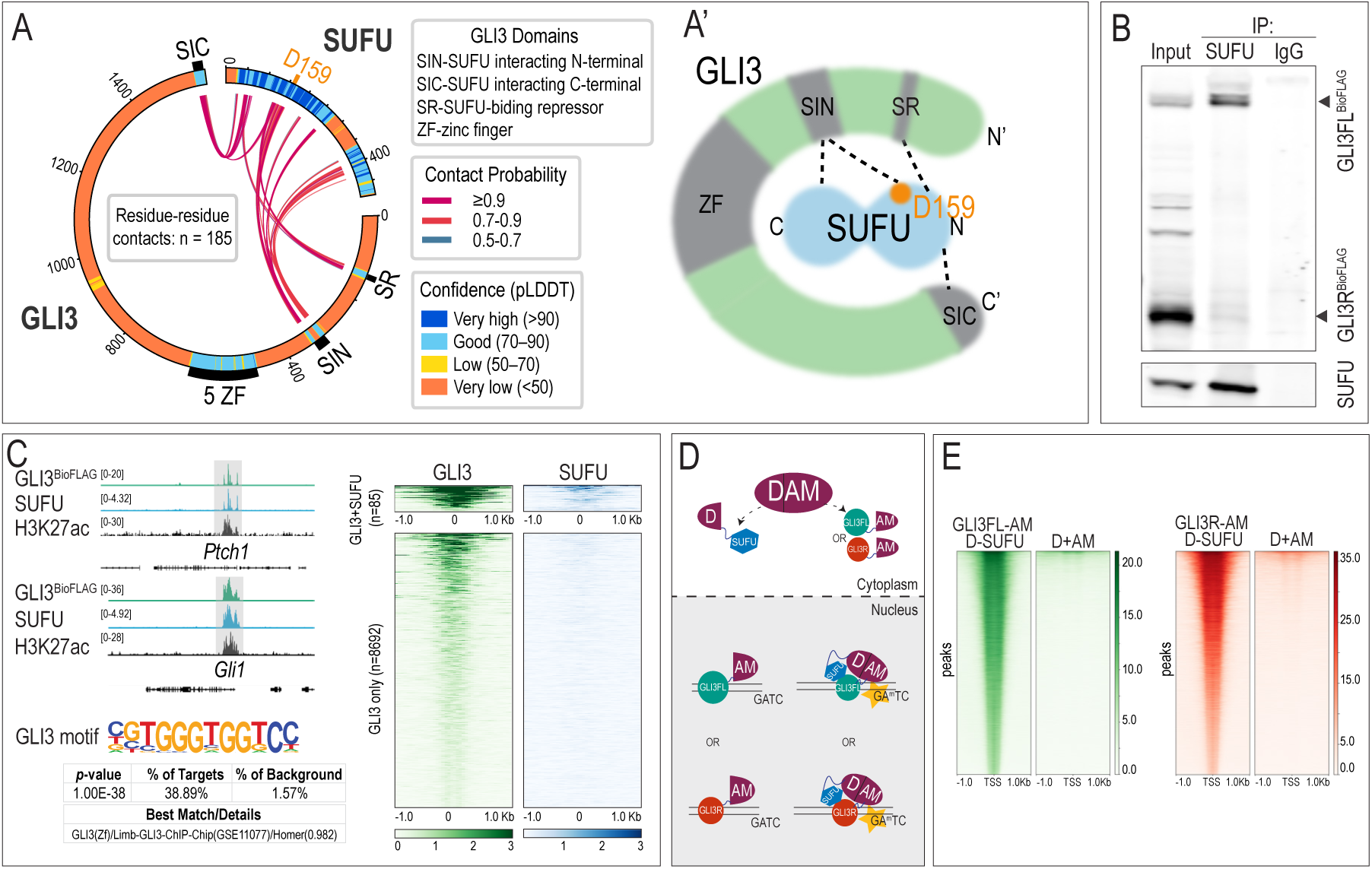
SUFU interacts with nuclear GLI3 and binds to DNA. (A) Chord diagram showing the pLDDT confidence of every residue, domain annotation (SIN: Sufu-interacting N-terminal domain; SIC: Sufu-interacting C-terminal domain; SR: SUFU-binding repressor domain; ZF: zinc finger DNA binding domain), and interchain contact. Chords linking SUFU residues to GLI3 residues (n=185) were colored based on contact probability. (A’) Simplified schematic of mammalian SUFU and GLI3 protein domains and interacting regions (dashed lines). Acidic residue D159 is located in the N-terminal lobe of SUFU and has contacts with the SIN domain of GLI3. (B) Co-immunoprecipitation of SUFU and GLI3 isoforms with anti-FLAG antibody in E11.5 *Gli3^BioFLAG/BioFLAG^* MNPs. Arrowheads indicate GLI3FL and GLI3R. (C) CUT&RUN showing both SUFU and GLI3 were enriched at *Ptch1* and *Gli1* loci that were also marked by histone H3K27ac. Heat maps for all GLI3 peaks (n = 8,777), partitioned by SUFU co-occupancy. SUFU co-occupies a subset of GLI3-bound sites with the highest signal (n = 85). Among the co-occupied regions, the GLI3 motif was the top ranked motif. (D) Schematic of Split DamID assay. D- or AM-domain of Dam DNA methylase was tagged to SUFU and either GLI3FL or GLI3R, respectively. (E) Heatmap of Split DamID results showing SUFU interaction with GLI3FL or GLI3R at GLI3 target genes.

To determine if SUFU contributed to GLI3-dependent transcriptional activity, we analyzed GLI3 and SUFU binding across the genome. CUT&RUN-seq using the *Gli3^BioFLAG^* murine line enabled analysis of endogenous GLI3 binding (34). GLI3 and SUFU CUT&RUN tracks were enriched at similar positions in the promoter regions of *Ptch1* and *Gli1*, coincident with the active promoter/enhancer mark H3K27ac (Fig. 1C) (35). Because SUFU does not bind DNA directly, its CUT&RUN signal was restricted to a few enrichment sites genome-wide (Fig. 1C and Supplementary Fig. 1C); nevertheless, 85 of 8777 GLI3 peaks (1%) coincided with SUFU peaks. A permutation test confirmed that this overlap far exceeded the overlap expected by chance (85 vs. ∼0.4 expected overlaps; Z-score 137.3; *p* < 1×10^-4^; Supplementary Fig. 1C). Further, a co-occupancy-partitioned heatmap showed that GLI3 signal was stronger at co-occupied (GLI3+SUFU) sites than at GLI3-only sites (n = 85 co-occupied peaks; Fig. 1C). HOMER *de novo* motif analysis of the co-occupied sites recovered the GLI3 motif as the top-ranked motif (*p* = 1×10^-38^; 38.9% of targets vs. 1.6% of background; Fig. 1C). Together, these data indicate that SUFU and GLI3 co-occupy the genome as part of a transcriptional complex.

To further test whether GLI3 and SUFU remain in complex on DNA, we performed Split DamID, a modified DamID assay designed to examine the functional relationships between interacting or juxtaposed TFs or coregulators on chromatin (36, 37). In this assay, the D and AM fragments of *E. coli* DNA adenine methyltransferase (Dam) are fused to the proteins of interest, such that their interaction or juxtaposition on DNA reconstitutes Dam and methylates adenine in nearby GATC sequences (Fig. 1D). D-SUFU with GLI3FL-AM or D-SUFU with GLI3R-AM were co-transfected into the O9-1 murine neural crest cell line (38) because many of the ciliopathic phenotypes observed are in areas derived from cranial neural crest cells. Construct expression was confirmed by western blotting (Supplementary Fig. 1D), and methylated DNA was sequenced. GLI3FL-AM:D-SUFU and GLI3R-AM:D-SUFU complexes marked a substantially greater number of GATC sequences than the D+AM control (Fig. 1E), indicating that SUFU was in complex with both GLI3FL and GLI3R at GLI3-binding regions. GO (gene ontology) analysis of these targets identified many genes associated with craniofacial phenotypes and abnormal craniofacial morphology (Table S2). To confirm that this interaction was reproducible, we repeated the assay in an independent cell line, MK4. SUFU-GLI3 complexes again marked GATC-flanked regions near GLI3 target genes, and GO analysis of these targets recovered terms associated with craniofacial development as well as abnormal nephron morphology, the latter consistent with the metanephric (kidney) identity of MK4 cells (Table S2). Because of technical differences between Split DamID (a broader 1-2 kb footprint and steric constraints from the fusion proteins) and CUT&RUN (precise, targeted cleavage), the two datasets correlated at the gene level rather than at individual binding sites. Nevertheless, these data support the conclusion that dissociation from SUFU is not a prerequisite for GLI3FL/GLI3R nuclear localization and indicate that SUFU actively contributes to GLI3-mediated transcriptional regulation.

### Disrupting SUFU-GLI interaction results in severe embryonic defects

Previous *in vitro* studies established the physical association of SUFU-GLI (30) through SIN and SIC domains and, in GLI2 and GLI3, a more recently identified SUFU-binding repressor (SR) domain (23, 29) (Fig. 1A-A′ and Supplementary Fig. 1B). A patch of seven acidic residues within the N-terminal lobe of SUFU is essential for this interaction (27, 30, 39). Within this acidic patch, the aspartic acid at the 159^th^ position (D159) is the only residue that is critical for SUFU-GLI interaction. Previous *in vitro* studies replacing D159 with an arginine residue (D159R) abolished SUFU binding to GLI (27, 29, 30). Consistent with this, AlphaFold3 modeling of the SUFU^D159R^-GLI3 complex predicted a markedly weakened interface (107 vs. 185 residue-residue contacts), with complete loss of contacts to the GLI3 SYGHLS motif within the SIN domain (52 contacts in wild type to none) (Table S1 and Supplementary Fig. 2A). Notably, SUFU^D159R^ adopted a more “open” conformation than SUFU^WT^, with its N- and C-terminal lobes splayed apart rather than clamped around the GLI3 SYGHLS motif, that also resulted in reduced inter-chain interactions and introduced intra-chain clashes (Supplementary Fig. 2B-C’). This “open” conformation resembles the unbound state of SUFU resolved by crystallography and small-angle X-ray scattering (39). Together, these data suggest that D159R disrupts SUFU-GLI3 engagement by preventing closure of the SUFU clamp around the SYGHLS motif within the GLI3 SIN domain. To test the biological consequence of D159R in SUFU-GLI3 interactions, we co-transfected MK4 cells, a murine metanephric mesenchymal line with high transfection efficiency (40), with GFP-tagged forms of either SUFU^WT^ or SUFU^D159R^ together with FLAG-tagged GLI3FL or GLI3R. GLI3 was then immunoprecipitated and its association with SUFU was analyzed by western blotting. The D159R mutation greatly reduced interactions between SUFU and both GLI3FL and GLI3R isoforms (Supplementary Fig. 2D, E; asterisks), indicating a critical role for D159 in SUFU-GLI3 interactions under wild-type, non-ciliopathic conditions. Moreover, whereas GFP-tagged SUFU^WT^ sequestered GLI2 and GLI3 in the cytoplasm, GFP-tagged SUFU^D159R^ relieved this sequestration, allowing both proteins to traffic into the nucleus (Supplementary Fig. 2F-H″). Thus, D159 is essential for SUFU-mediated cytoplasmic sequestration of GLIs, and GLI2 and GLI3 were affected similarly by the D159R mutation. Because GLI3 is the predominant GLI family member in craniofacial neural crest-derived tissues, where both its activator and repressor functions are required across the MNP and FNP (15), and because the SUFU-interacting domains (SIN, SIC and SR) are conserved among GLI proteins (Supplementary Fig. 1B), we focused subsequent *in vivo* analyses on GLI3.

To determine the developmental consequences of impaired SUFU-GLI3 interactions, we generated mice carrying the *Sufu^D159R^* mutation using CRISPR-Cas9 editing to convert GAC to AGA in the 5′ region of exon 4 (Asp to Arg; Supplementary Fig. 3A). Heterozygous *Sufu^D159R/+^* embryos were viable and did not display gross morphological phenotypes (Fig. 2A-B’). In contrast, homozygous *Sufu^D159R/D159R^* mutant embryos exhibited severe developmental defects resembling *Sufu^-/-^* embryos (41), including an open neural tube, under-developed craniofacial complex, and early embryonic lethality (Fig. 2C-D’). A complementation test with the compound *Sufu^D159R/-^* mutant produced similar abnormalities (Fig. 2E-E’). Despite similar phenotypic presentations, RNA-seq analysis on E9.5 embryonic heads (at least 3 replicates per genotype) revealed significant transcriptional differences between the two genotypes. Principal component analysis showed *Sufu^-/-^* embryos had the most pronounced transcriptional changes relative to wild-type, whereas *Sufu^D159R/D159R^* mutants exhibited a more moderate shift (Fig. 2F). This result, together with the predicted 107 remaining contacts between SUFU^D159R^ and GLI3 (Supplementary Fig. 2A and Table S1), suggested that the *Sufu^D159R^* mutation altered SUFU-GLI binding, rather than eliminating all physical contact. This altered binding allows for isolating the specific contribution of the SUFU-GLI binding interface to the observed phenotypes. Gene expression analysis identified genes with a significant, greater than 2-fold, change between *Sufu^-/-^* and wild-type, and between Sufu*^D159R/D159R^* and wild-type (Table S3 and Table S4). Notably, some genes in the *Homeobox* and *Fox* (Forkhead box) families were differentially expressed in the same direction in both *Sufu^-/-^* and Sufu*^D159R/D159R^* mutants. Among HH pathway genes, *Shh, Ptch1, Gli3,* but not *Gli1* were differentially expressed in both mutants (Table S4). The lack of *Gli1* dysregulation was confirmed by RT-qPCR (Supplementary Fig. 3B) and was consistent with published *Sufu null* data, in which *Gli1* expression was not clearly altered in the head region despite ventralization of the neural tube (41). Thus, despite similar phenotypic presentation, *Sufu^D159R^*^/*D159R*^ had a distinct transcriptional profile from *Sufu*^-/-^.

**Figure 2.**
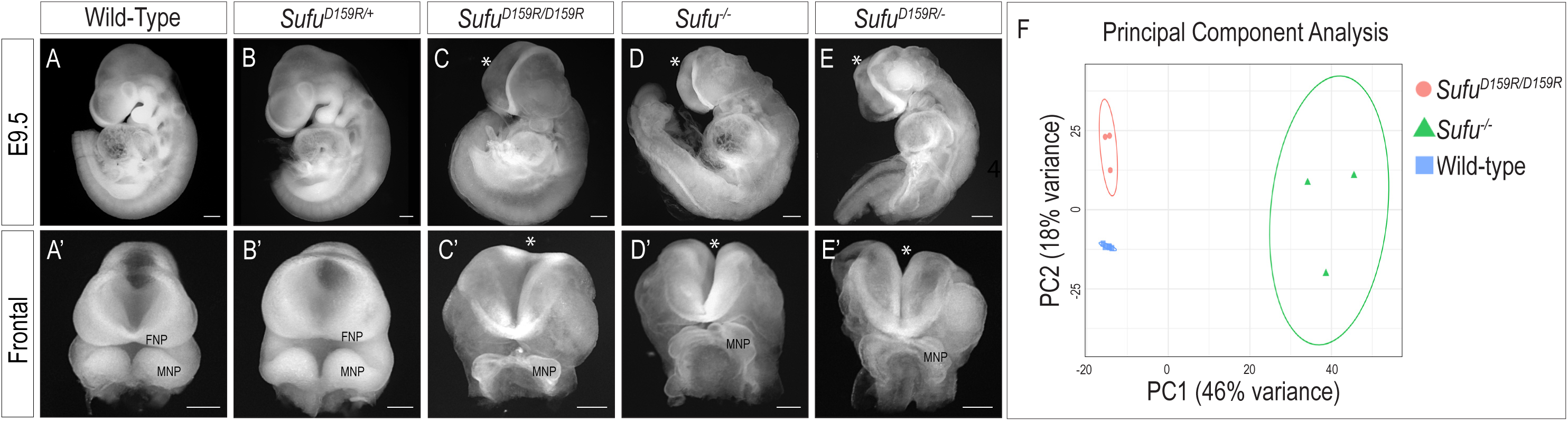
Disrupting SUFU-GLI3 interactions results in severe embryonic defects. (A-E) Lateral and (A’-E’) frontal views of E9.5 wild-type, *Sufu^D159R/+^* heterozygous, *Sufu^D159R/D159R^* homozygous, *Sufu^-/-^* homozygous, and *Sufu^D159R/-^* compound mutant embryos. Both homozygous and compound mutants exhibit loss of facial prominences, open neural tube (*), and early embryonic lethality (n ≥ 3 for all genotypes). (F) Principal component analysis displaying the transcriptomic profiles of wild-type, homozygous *Sufu^D159R/D159R^* and *Sufu^-/-^* mutants. Each point represents an individual biological replicate, and each ellipse denotes 95% confidence intervals for each genotype group. PC1 and PC2 account for 46% and 18% of the total variance, respectively. MNP; mandibular prominence. FNP; frontonasal prominence. Scale bar = 200 μm.

### SUFU D159 residue modulates SUFU-GLI interactions through cilium-dependent and cilium-independent mechanisms

SUFU^D159R^ altered binding with GLI3 *in vitro* (Supplementary Fig. 2). To determine whether this mutation affects GLI3 binding *in vivo*, we performed immunoprecipitation with a SUFU antibody on lysates from E9.5 mouse heads. Interactions between SUFU and GLI3 proteins were slightly reduced in *Sufu^D159R/+^* embryos and markedly lost in *Sufu^D159R/D159R^* mutant embryos (Fig. 3A asterisks), indicating that SUFU^D159R^ abrogated SUFU-GLI3 interactions and affected protein stability of GLI3 *in vivo* under wild-type, non-ciliopathic conditions. Given the previous work reporting an epistatic relationship between SUFU and the cilium, we next asked how SUFU^D159R^ affected SUFU-GLI3 interactions in the absence of cilia. One allele of *Sufu^D159R^* was crossed onto the *Kif3a^fl/fl^;Wnt1Cre;Gli3^BioFLAG/+^* background and co-immunoprecipitation for SUFU-GLI3 complexes from E11.5 MNP tissue lysates of control *Kif3a^fl/fl^;Gli3^BioFLAG/+^*, *Kif3a^fl/fl^;Wnt1Cre;Gli3^BioFLAG/+^*, and *Kif3a^fl/fl^;Wnt1Cre;Sufu^D159R/+^;Gli3^BioFLAG/+^* embryos was performed. Notably, SUFU^D159R^ still pulled down GLI3 isoforms on the ciliopathic background (Fig. 3B). This residual interaction likely occurred through the SIC domain and an internal region that hinders coactivator binding, as previously reported (27). These results uncovered two distinct SUFU-GLI binding modes: a cilium-dependent binding mode that involves the D159 contact interface which is the predominant interaction when cilia are functional and that is also required for GLI protein stability (Fig. 3A’); and a cilium-independent binding mode that does not rely on D159, using an alternative interaction interface that becomes the predominant interaction when cilia are absent (Fig. 3B’).

**Figure 3.**
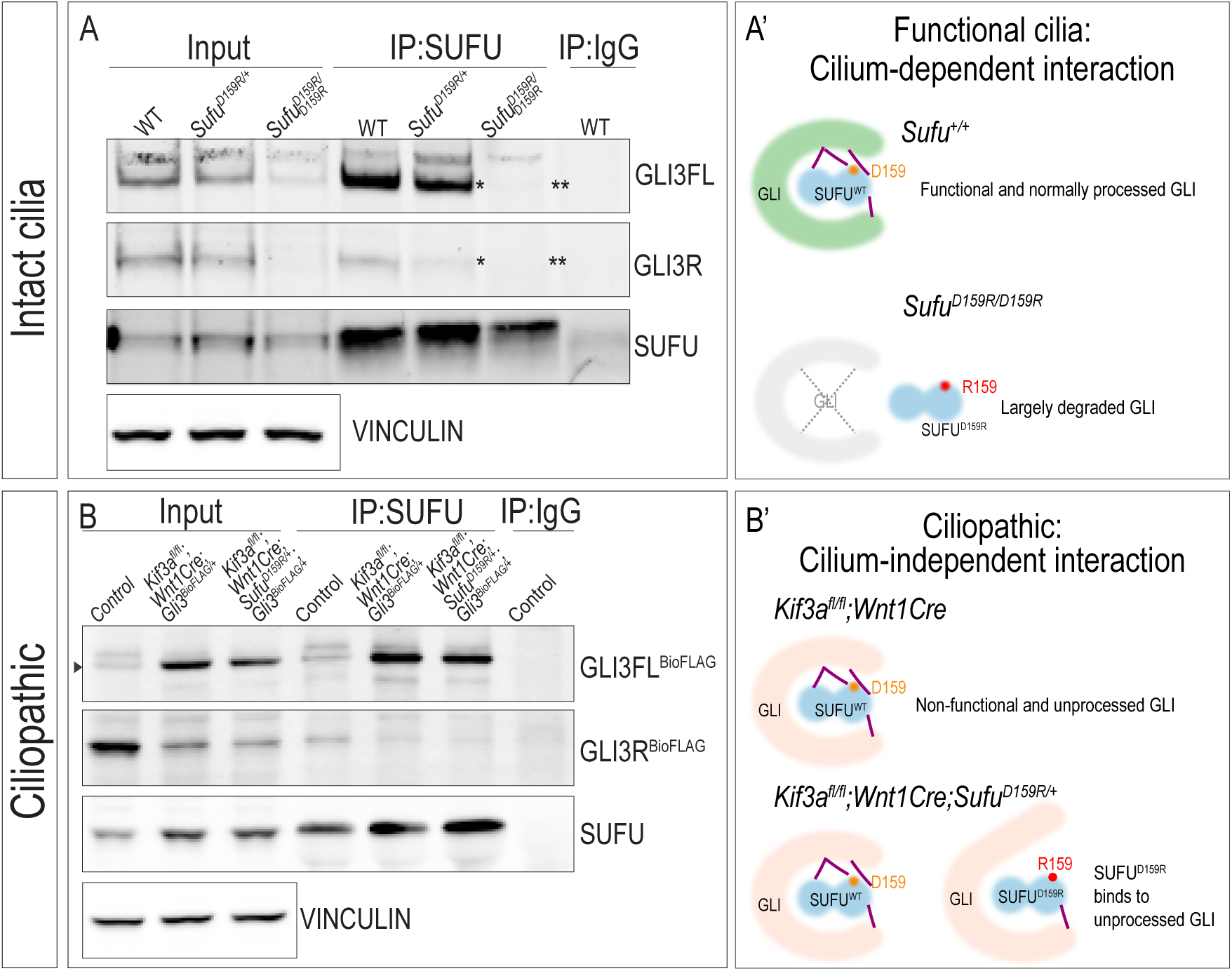
SUFU interacts with GLI3 isoforms via cilia-dependent and cilia-independent interactions. (A) Co-immunoprecipitation of GLI3 with an anti-SUFU antibody in E9.5 wild-type (n=13), *Sufu^D159R/+^* (n=24), or *Sufu^D159R/D159R^* (n=10) whole embryo lysate. Asterisks denote reduced (*) or undetectable (**) GLI3 protein. (A’) Schematic of SUFU-GLI3 interactions in the presence of functional cilia. SUFU interacts with GLI3, GLI3 is trafficked through the cilia and post-tranlationally processed into a functional GLI3 protein. *Sufu^D159R/D159R^* mutation abrogates SUFU-GLI3 interactions, resulting in largely degraded GLI3. (B) Co-Immunoprecipitation of GLI3 with an anti-SUFU antibody in pooled MNPs from E11.5 *Kif3a^fl/fl^;Gli3^BioFLAG/+^* (control, n=22), *Kif3a^fl/fl^;Wnt1Cre;Gli3^BioFLAG/+^* (n=18) and *Kif3a^fl/fl^;Wnt1Cre;Sufu^D159R/+^*;*Gli3^BioFLAG/+^* (n=15) embryos. Under ciliopathic conditions, introduction of *Sufu^D159R/+^* does not abrogate SUFU-GLI3 interactions. (B’) Schematic of SUFU-GLI3 interactions under ciliopathic conditions. In the absence of cilia, SUFU interacts with GLI3, but GLI3 does not traffic through the cilia and is not post-translationally processed, resulting in a non-functional GLI3 protein. Introduction of the *D159R* allele allows for maintained SUFU binding to the GLI3-SIC domain, resulting in a partially active GLI3.

### Introduction of the *Sufu^D159R^* allele ameliorates ciliopathic craniofacial phenotypes

Typical of several conditional ciliopathic mutants, *Kif3a^fl/fl^;Wnt1Cre* embryos present with craniofacial phenotypes including midfacial widening, bifurcation of the nasal septum, bilateral cleft palate, micrognathia and hypoglossia/aglossia (15, 16). These phenotypes are both intriguing and problematic as the midfacial phenotypes are associated with a net gain of HH signaling due to loss of GLIR activity, whereas the lower jaw phenotypes are associated with a net loss of HH signaling due to a loss of GLIA activity. Since ciliopathic mutants presented with excessive SUFU-GLI3 complexes, we tested whether introducing *Sufu^+/-^* or *Sufu^D159R^* could rescue midfacial and lower facial phenotypes in *Kif3a^fl/fl^;Wnt1Cre* mice. Both *Kif3a^fl/fl^;Wnt1Cre;Sufu^+/-^* and *Kif3a^fl/fl^;Wnt1Cre;Sufu^D159R/+^* presented partially rescued midfacial phenotypes (e.g., single medial cleft and single cartilaginous nasal septum) (Fig. 4A-C’’’). Quantification of the ratio of the distance between nasal pits and distance between eyes showed introduction of both *Sufu^+/-^* and *Sufu^D159R^* alleles were able to partially restore the midfacial widening phenotype of *Kif3a^fl/fl^;Wnt1Cre* to a similar degree (Fig. 4D). In the lower face, however, only *Kif3a^fl/fl^;Wnt1Cre;Sufu^D159R/+^* embryos restored the projection of the mandible and area of the tongue to levels not significantly different from wild-type embryos (Fig. 4E-I). In addition to facial phenotypes, epithelial specializations (e.g., in tooth development) were also partially rescued in *Kif3a^fl/fl^;Wnt1Cre;Sufu^D159R/+^* embryos (Supplementary Fig. 4A-B”). Together, these results suggested that while the introduction of both *Sufu^+/-^* and *Sufu^D159R/+^* rescued midfacial development in the absence of cilia, only *Sufu^D159R/+^* rescued mandibular and glossal development to a degree that was not significantly different than wild-type.

**Figure 4.**
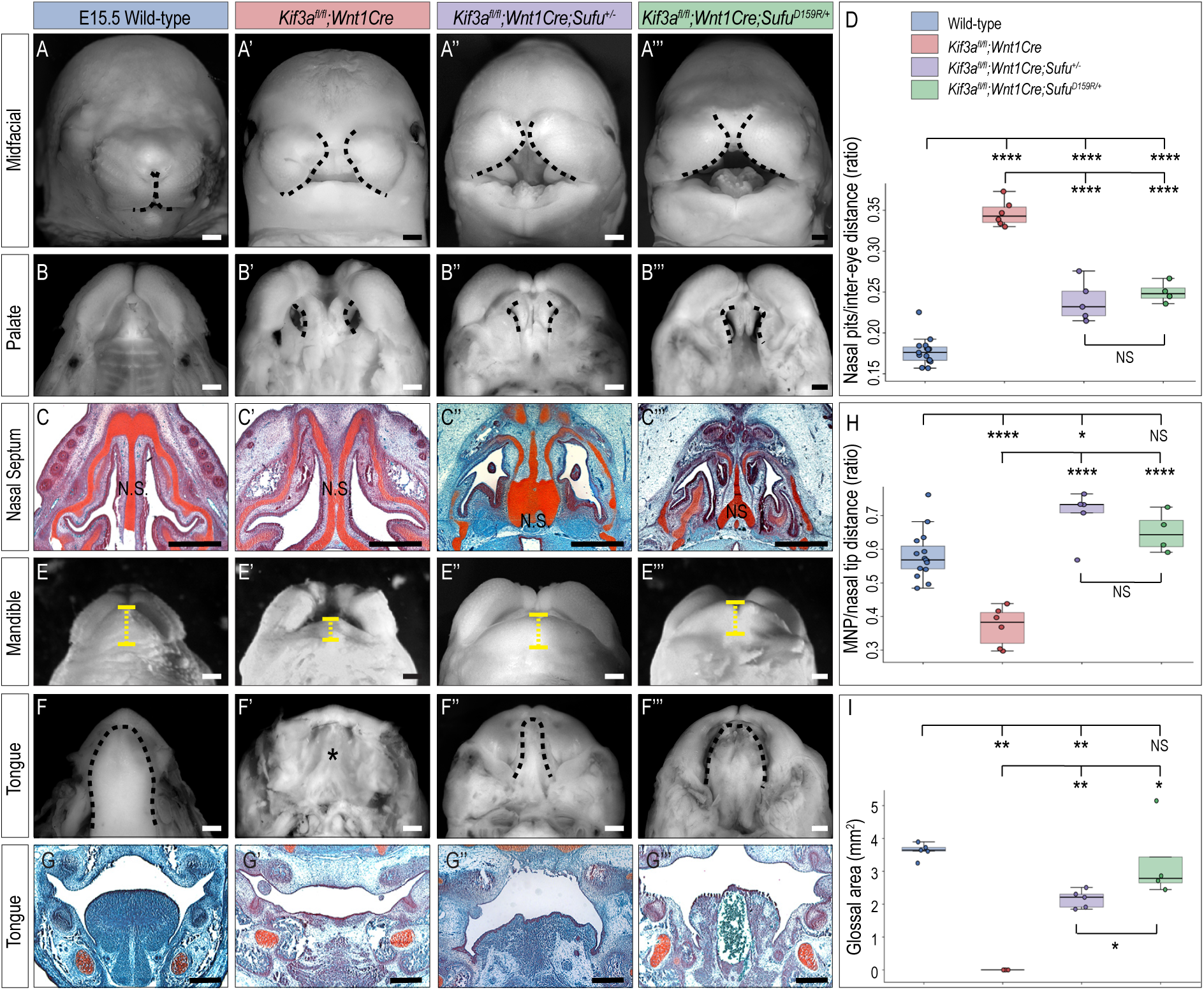
Introduction of *Sufu^D159R^* or *Sufu^null^* alleles partially rescue ciliopathic phenotypes. (A-C’’’) Midfacial phenotypes in E15.5 wild-type, *Kif3a^fl/fl^;Wnt1Cre, Kif3a^fl/fl^;Wnt1Cre;Sufu^+/-^*, and *Kif3a^fl/fl^; Wnt1Cre;Sufu^D159R/+^* embryos including (A-A’’’) midfacial widening (dotted black line), (B-B’’’) cleft palate (dotted black line), and (C-C’’’) bifurcation of the nasal septum (N.S.). (D) Quantitative measurement of midfacial widening via the relative nasal pits/inter-eye distance reveals that both *Sufu^+/-^* and *Sufu^D159R/+^* alleles significantly restore midfacial width. (E-G’’’) Mandibular phenotypes in E15.5 wild-type, *Kif3a^fl/fl^;Wnt1Cre, Kif3a^fl/fl^;Wnt1Cre;Sufu^+/-^*, and *Kif3a^fl/fl^;Wnt1Cre;Sufu^D159R/+^* embryos including (E-E’’’) micrognathia (dotted yellow line) and (F-G’’’) aglossia (compare dotted black line in F and asterisk in F’). (H,I) Quantitative measurement of mandibular projection and glossal area reveals that introduction of the *Sufu^D159R/+^* allele onto the ciliopathic background results in the most complete rescue of mandibular/ glossal phenotypes. (N ≥ 3 for all genotypes). Scale bar = 500 μm.

### Introduction of *Sufu^D159R^* restores ability of GLI3 to bind to DNA and transcribe targets in the absence of cilia

The rescue of ciliopathic phenotypes associated with both the introduction of *Sufu^+/-^* and *Sufu^D159R/+^* alleles was intriguing, yet the mechanistic basis for the rescue was unclear. Previous reports have shown that SUFU regulated GLI-directed transcriptional activity (22, 29). Thus, we examined how addition of *Sufu^+/-^* or *Sufu^D159R/+^* in E11.5 *Kif3a^fl/fl^;Wnt1Cre;Sufu^+/-^* and *Kif3a^fl/fl^;Wnt1Cre;Sufu^D159R/+^* FNPs and MNPs affected gene expression, compared to wild-type and *Kif3a^fl/fl^;Wnt1Cre*. Using a threshold of adjusted *p* < 0.05 and absolute log2 fold-change > 1, we identified 22 genes in the FNP and 39 in the MNP that were dysregulated in *Kif3a^fl/fl^;Wnt1Cre* compared to wild-type (Fig. 5A and Table S5). We then defined a gene as restored when the addition of either *Sufu^+/-^* or *Sufu^D159R/+^* significantly shifted it in the direction opposite to the mutant defect (*Kif3a^fl/fl^;Wnt1Cre;Sufu^+/-^* or *Kif3a^fl/fl^;Wnt1Cre;Sufu^D159R/+^* versus *Kif3a^fl/fl^;Wnt1Cre*, adjusted *p* < 0.05 with opposite direction of log2 fold-change). By this criterion, 10 of the 22 FNP genes and 14 of the 39 MNP genes were significantly restored by at least one *Sufu*-variant allele (Fig. 5B). In the FNP, restoration was largely shared between the two *Sufu* allele variants tested (7 genes restored by both, 3 by *Sufu^D159R^* alone, none by *Sufu^+/-^* alone), whereas in the MNP 9 genes were restored by both, 4 by *Sufu^D159R^* alone, and 1 by *Sufu^+/-^* alone. To quantify how completely each allele corrected the mutant transcriptome, we calculated the fraction of the *Kif3a^fl/fl^;Wnt1Cre* dysregulation returned toward wild-type by each allele for every dysregulated gene (Fig. 5A). In the FNP both alleles produced near-complete correction, with no significant difference between them (median fraction corrected 0.98 for *Sufu^D159R^* versus 0.95 for *Sufu^+/-^*; two-sided exact Wilcoxon signed-rank test, *p* = 0.085, n = 22). In the MNP, however, correction was significantly greater for *Sufu^D159R^* than for *Sufu^+/-^* (median 0.37 versus 0.18; *p* = 4.40 × 10^-5^, n = 39), indicating that the *Sufu^D159R^* allele more effectively restored the MNP transcriptome. These data were consistent with phenotypic presentation and degree of rescue of *Kif3a^fl/fl^;Wnt1Cre;Sufu^+/-^* and *Kif3a^fl/fl^;Wnt1Cre;Sufu^D159R/+^* embryos. Examination of specific differentially expressed genes revealed both up- and down-regulated genes in *Kif3a^fl/fl^;Wnt1Cre* embryos compared to wild-type in FNPs and MNPs, suggesting that loss of cilia predominantly impaired GLIFL activator (GLIA) function; however, a smaller proportion of genes regulated by GLIR were also restored (Fig. 5B and Table S5). GO analysis of the differentially expressed genes particularly in MNPs between *Kif3a^fl/fl^;Wnt1Cre* and wild-type, and between *Kif3a^fl/fl^;Wnt1Cre;Sufu^D159R/+^* and *Kif3a^fl/fl^;Wnt1Cre* revealed genes involved in abnormal craniofacial/palatal development and neural crest differentiation (Supplementary Fig. 5A-B’). Interestingly, members of the *Fox* TF family were significantly down-regulated in *Kif3a^fl/fl^;Wnt1Cre* and restored in *Kif3a^fl/fl^;Wnt1Cre;Sufu^D159R/+^* embryos (Fig. 5B, C and Supplementary Fig. 5A-A’), whereas members of the *Homeobox* and *T-box* TF families were up-regulated in *Kif3a^fl/fl^;Wnt1Cre* and restored in *Kif3a^fl/fl^;Wnt1Cre;Sufu^D159R/+^* embryos (Supplementary Fig. 5B-B’). CollecTRI analysis (42) revealed several of the differentially expressed genes, including multiple *Fox* TFs, were GLI targets (Supplementary Fig. 5C, labeled genes). Collectively, the transcriptomic results suggested that introducing a single allele of *Sufu^D159R^* was a more effective rescue, both phenotypically and molecularly, than *Sufu^+/-^*, and was sufficient to restore the expression of key GLI targets critical for craniofacial development. Thus, we continued our analysis focusing on the mechanism of *Sufu^D159R^*.

**Figure 5.**
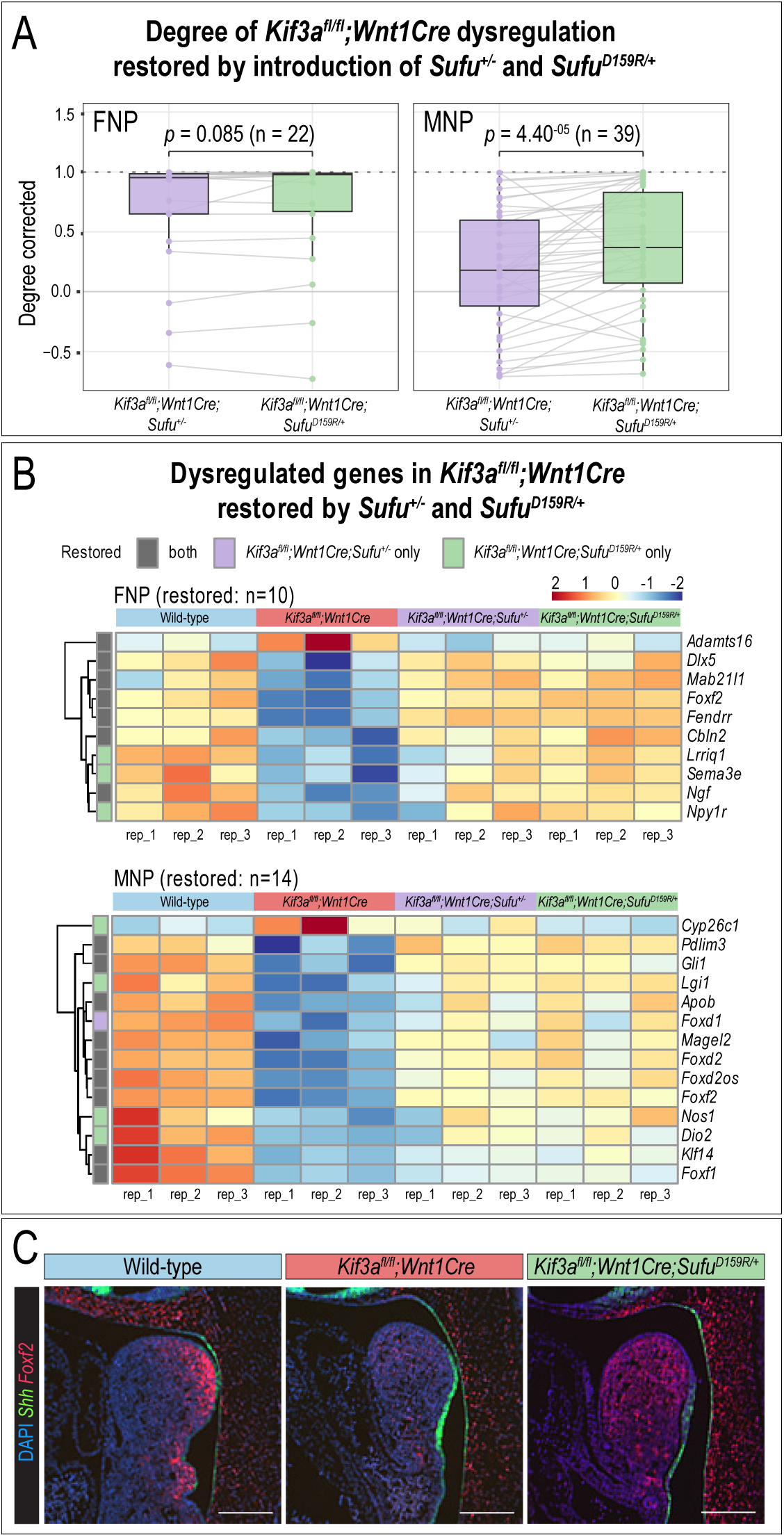
Introduction of *Sufu^D159R/+^* molecularly restores gene expression in ciliopathic FNPs and MNPs. (A) Degree of restored gene expression in the FNP (n = 22) or MNP (n = 39) between *Kif3a^fl/fl^;Wnt1Cre* and either *Kif3a^fl/fl^;Wnt1Cre;Sufu^+/-^* or *Kif3a^fl/fl^;Wnt1Cre;Sufu^D159R/+^*; higher values indicate greater restoration toward wild-type. *Sufu^D159R/+^* and *Sufu^+/-^* equally restored expression in the FNP, but *Sufu^D159R/+^* provides a significantly improved molecular rescue in the MNP (two-sided exact Wilcoxon signed-rank test; FNP *p* = 0.085; MNP *p* = 4.40 × 10^-5^). (B) Heatmaps of dysregulated genes in *Kif3a^fl/fl^;Wnt1Cre* mutants significantly restored by *Sufu^+/-^* and/or *Sufu^D159R/+^* (FNP, n = 10; MNP, n = 14). Far left column indicates whether each gene was restored by both alleles (grey), *Sufu^+/-^* only (purple), or *Sufu^D159R/+^* only (green) (n=3 per genotype). (C) RNAscope for *Shh* and *Foxf2* in E10.5 MNPs (sagittal view), showing restoration of *Foxf2* by the *Sufu^D159R/+^* allele. Scale bar = 200 µm; n = 3.

Based on observed transcriptional changes and previous reports that SUFU-GLI association was required for GLI to bind to DNA (14, 22), we performed CUT&RUN on MNPs (43) to measure endogenous GLI3 binding (34, 44) to genomic DNA in control, *Kif3a^fl/fl^;Wnt1Cre;Gli3^BioFLAG/+^,* and *Kif3a^fl/fl^;Wnt1Cre;Sufu^D159R/+^;Gli3^BioFLAG/+^* embryos. In control embryos, GLI3 showed enrichment in 8777 peaks (Table S6). Loss of cilia in *Kif3a^fl/fl^;Wnt1Cre;Gli3^BioFLAG/+^* embryos significantly reduced GLI3 binding to only 259 peaks, suggesting that cilia were essential for GLI3 binding to DNA. Introduction of *Sufu^D159R^* restored GLI3 binding at 2782 peaks (Fig. 6A; Table S6), a 32% (∼11-fold) restoration of DNA-binding, including at established GLI3 targets essential for HH pathway transduction (Fig. 6B). CUT&RUN with a SUFU antibody revealed that SUFU enrichment mirrored that of GLI3, with enrichment at *Ptch1* and *Gli1* in control that was reduced in *Kif3a^fl/fl^;Wnt1Cre;Gli3^BioFLAG/+^* and restored in *Kif3a^fl/fl^;Wnt1Cre;Sufu^D159R/+^;Gli3^BioFLAG/+^* embryos (Fig. 6B), supporting a role for SUFU in GLI3 transcriptional complexes. We note that the sequencing depth achieved in this study using *Gli3^BioFLAG/+^* may limit sensitivity for weaker or more transient GLI3 binding events, particularly at distal enhancers or low-occupancy sites. Nevertheless, these data successfully identified established GLI3 target loci, and the genome-wide peak set showed strong concordance with our independently generated SUFU CUT&RUN data (Fig. 6B).

**Figure 6.**
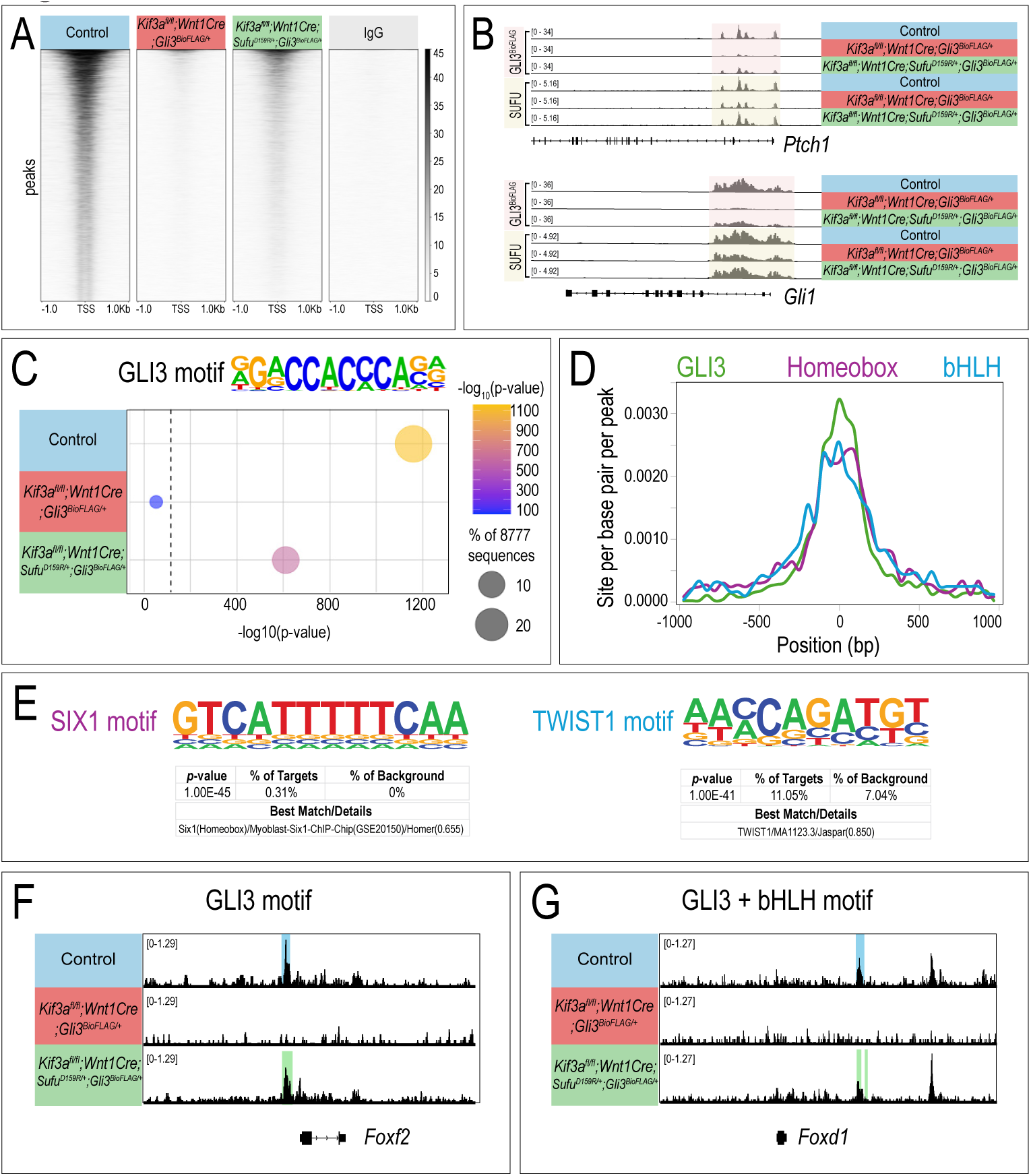
Introduction of *Sufu^D159R/+^* restores DNA-binding ability of GLI3. (A) Heatmap of CUT&RUN from E11.5 *Kif3a^fl/fl^;Gli3^BioFLAG/+^*(control), *Kif3a^fl/fl^;Wnt1Cre;Gli3^BioFLAG/+^*, and *Kif3a^fl/fl^;Wnt1Cre;Sufu^D159R/+^;Gli3^BioFLAG/+^;* MNPs. (B) Genome tracks of GLI3 (FLAG) and SUFU CUT&RUN at *Ptch1* and *Gli1* loci. (C) Motif analysis showing the GLI3 motif is enriched in GLI3-bound regions from control and *Kif3a^fl/fl^;Wnt1Cre;Sufu^D159R/+^;Gli3^BioFLAG/+^* MNPs but absent in *Kif3a^fl/fl^; Wnt1Cre;Gli3^BioFLAG/+^* MNPs (dashed line indicates statistical significance threshold). (D) Homeobox and bHLH TF motifs were frequently located near the GLI3 motif. (E) HOMER *de novo* motif analysis showing examples of Homeobox (SIX1) and bHLH (TWIST1) motifs. (F-G) Examples of GLI3 and bHLH motifs in GLI3 peaks near *Foxf2* and *Foxd1* promoter regions that were lost in *Kif3a^fl/fl^;Wnt1Cre; Gli3^BioFLAG/+^* but restored in *Kif3a^fl/fl^;Wnt1Cre;Sufu^D159R/+^;Gli3^BioFLAG/+^* MNPs.

We next examined the specific loci of restored binding in *Kif3a^fl/fl^;Wnt1Cre;Sufu^D159R/+^;Gli3^BioFLAG/+^* embryos. Known motif analysis confirmed the presence of conserved GLI motifs in wild-type MNPs and revealed restored binding at these motifs in *Kif3a^fl/fl^;Wnt1Cre;Sufu^D159R/+^;Gli3^BioFLAG/+^* MNPs at known targets (Fig. 6C). GO analysis of the genes at loci where GLI3 binding was lost in *Kif3a^fl/fl^;Wnt1Cre;Gli3^BioFLAG/+^* and restored in *Kif3a^fl/fl^;Wnt1Cre;Sufu^D159R/+^;Gli3^BioFLAG/+^* MNPs identified genes involved in transcription factor binding, transcription coregulator activity, chromatin binding, and craniofacial morphology (Table S6). Our work, and that of others, has reported that GLI TFs work with cofactors to execute tissue-specific gene regulatory networks (45, 46). Interestingly, by HOMER *de novo* motif analysis, Homeobox and bHLH motifs were identified within GLI3 peaks (Fig. 6D-E). These peaks are associated with genes restored in *Kif3a^fl/fl^;Wnt1Cre;Sufu^D159R/+^;Gli3^BioFLAG/+^* samples, including *Fox* TFs (Fig. 6F-G) which were previously shown to be misexpressed in *Kif3a^fl/fl^;Wnt1Cre;Gli3^BioFLAG/+^*, and restored in *Kif3a^fl/fl^;Wnt1Cre;Sufu^D159R/+^;Gli3^BioFLAG/+^* MNPs (Fig. 5B and Supplementary Fig. 5C). These findings suggest that alteration of SUFU-GLI3 complexes under ciliopathic conditions bypass the requirement of cilia in regulating GLI3 binding to DNA and binding to transcriptional cofactors.

It is possible that the rescue of ciliopathic phenotypes resulted from alternative mechanisms such as restoration of cilium extension, restoration of the GLI3FL/GLI3R ratio, and/or SUFU-mediated cytoplasmic sequestration of GLI3. To investigate these possibilities, we first examined cilium extension in mouse embryonic fibroblasts (MEFs) isolated from control *Kif3a^fl/fl^;Gli3^BioFLAG/+^*, *Kif3a^fl/fl^;Wnt1Cre;Gli3^BioFLAG/+^*, and *Kif3a^fl/fl^;Wnt1Cre;Sufu^D159R/+^;Gli3^BioFLAG/+^* facial prominences. Ciliary extension was reduced in both *Kif3a^fl/fl^;Wnt1Cre;Gli3^BioFLAG/+^* and *Kif3a^fl/fl^;Wnt1Cre;Sufu^D159R/+^;Gli3^BioFLAG/+^* relative to control MEFs (Supplementary Fig. 6A-D), indicating that the phenotypic rescue was not due to restoration of ciliary extension. Second, the skewed GLI3FL/GLI3R ratio was similar in *Kif3a^fl/fl^;Wnt1Cre;Gli3^BioFLAG/+^* and *Kif3a^fl/fl^;Wnt1Cre;Sufu^D159R/+^;Gli3^BioFLAG/+^* MNPs in both the input and the SUFU immunoprecipitants (Fig. 3B, input, IP), indicating that the rescue was not associated with restoration of the GLI3FL/GLI3R ratio. Finally, we asked whether SUFU-mediated cytoplasmic sequestration of GLI3 could account for the rescue by measuring nuclear GLI3FL and GLI3R with nuclear fractionation (Supplementary Fig. 6E). Both *Kif3a^fl/fl^;Wnt1Cre and Kif3a^fl/fl^;Wnt1Cre;Sufu^D159R/+^* MNPs had more nuclear GLI3FL and less nuclear GLI3R when compared with the control MNPs, indicating that SUFU^D159R^ did not alter cytoplasmic sequestration or nuclear localization of GLI isoforms in the absence of cilia, in contrast to its cellular functions in the presence of cilia (Supplementary Fig. 2F-H’’). Thus, the impressive rescue of ciliopathic phenotypes by addition of one allele of *Sufu^D159R^* could not be explained by restoration of cilium extension, restoration of the GLI3FL/GLI3R ratio, or cytoplasmic sequestration of GLI3. Rather, the restoration of GLI3 binding to DNA and GLI3 transcriptional activity were the most plausible cause of this rescue.

### Cofactor binding and post-translational modifications are restored with the addition of SUFU^D159R^

DNA binding and transcriptional activity of SUFU-GLI3 complexes were restored in *Kif3a^fl/fl^;Wnt1Cre;Sufu^D159R/+^* embryos, potentially through restoring interactions with transcriptional cofactors. To investigate this hypothesis, we analyzed chromatin-associated GLI3 protein complexes using <u>R</u>apid Immunoprecipitation <u>M</u>ass Spectrometry of <u>E</u>ndogenous proteins (RIME) (Fig. 7A) (47). RIME analysis was performed on E11.5 MNPs using *Gli3^BioFLAG/+^*, an endogenously FLAG-tagged *Gli3* allele to identify interacting proteins and to perform differential protein abundance analysis comparing *Gli3^BioFLAG/+^* controls versus *Kif3a^fl/fl^;Wnt1Cre;Gli3^BioFLAG/+^*, *Kif3a^fl/fl^;Gli3^BioFLAG/+^* controls versus *Kif3a^fl/fl^;Wnt1Cre;Sufu^D159R/+^;Gli3^BioFLAG/+^*, and *Kif3a^fl/fl^;Wnt1Cre;Gli3^BioFLAG/+^* versus *Kif3a^fl/fl^;Wnt1Cre;Sufu^D159R/+^*;*Gli3^BioFLAG/+^* (Table S7). First, we examined recovery of the target protein, GLI3, from both cytoplasmic and nuclear fractions relative to control, and observed enrichment at the peptide and whole-protein level (Table S7). Quantification of GLI3 protein abundance in the cytoplasm and nucleus of *Kif3a^fl/fl^;Gli3^BioFLAG/+^* control, *Kif3a^fl/fl^;Wnt1Cre;Gli3^BioFLAG/+^*, and *Kif3a^fl/fl^;Wnt1Cre;Sufu^D159R/+^;Gli3^BioFLAG/+^* samples indicated that GLI3 levels were elevated in both cytoplasmic and nuclear compartments in *Kif3a^fl/fl^;Wnt1Cre;Gli3^BioFLAG/+^* and *Kif3a^fl/fl^;Wnt1Cre;Sufu^D159R/+^*;*Gli3^BioFLAG/+^* MNPs compared to *Kif3a^fl/fl^;Gli3^BioFLAG/+^* controls (Fig. 7B, C). This increase in GLI3 abundance was consistent with the accumulation of unprocessed GLI3FL previously shown by western blot (Supplementary Fig. 6E). Interestingly, in *Kif3a^fl/fl^;Wnt1Cre;Sufu^D159R/+^*;*Gli3^BioFLAG/+^* samples, the cytoplasmic abundance of GLI3 was reduced in the presence of SUFU^D159R^ compared to *Kif3a^fl/fl^;Wnt1Cre;Gli3^BioFLAG/+^* (Fig. 7B). This reduction was not a result of impaired cytoplasmic sequestration, as nuclear GLI3 levels were not increased in the *Kif3a^fl/fl^;Wnt1Cre;Sufu^D159R/+^*;*Gli3^BioFLAG/+^* (Fig. 7C). Instead, the decreased cytoplasmic GLI3 may reflect enhanced protein degradation. We next examined the chromatin-associated GLI3 interactome to determine whether transcriptional cofactor binding was altered across conditions. Differential protein abundance analysis across all three pairwise comparisons (Table S7) revealed that a number of known transcriptional regulators were differentially associated with GLI3 in *Kif3a^fl/fl^;Wnt1Cre;Gli3^BioFLAG/+^* versus *Gli3^BioFLAG/+^* control and *Kif3a^fl/fl^;Wnt1Cre;Sufu^D159R/+^*;*Gli3^BioFLAG/+^* embryos. Among these, several nuclear FOX/GLI3 complexes were reduced in the *Kif3a^fl/fl^;Wnt1Cre;Gli3^BioFLAG/+^* and restored in the *Kif3a^fl/fl^;Wnt1Cre;Sufu^D159R/+^*;*Gli3^BioFLAG/+^* samples (Fig. 7D). This finding was particularly intriguing because several *Fox* TFs showed restored expression in *Kif3a^fl/fl^;Wnt1Cre;Sufu^D159R/+^* samples and restored GLI3 binding at their promoter region (Fig. 5B; Fig. 6F-G). These results suggested a positive feedback loop whereby GLI3 promotes transcription of genes that function as GLI cofactors. Thus, RIME analysis provided proteomic evidence that introduction of the *Sufu^D159R^* allele restored GLI3 interactions with transcriptional cofactors that were disrupted in the ciliopathic condition, and that these restored interactions included proteins with established roles in craniofacial development (Table S7).

**Figure 7.**
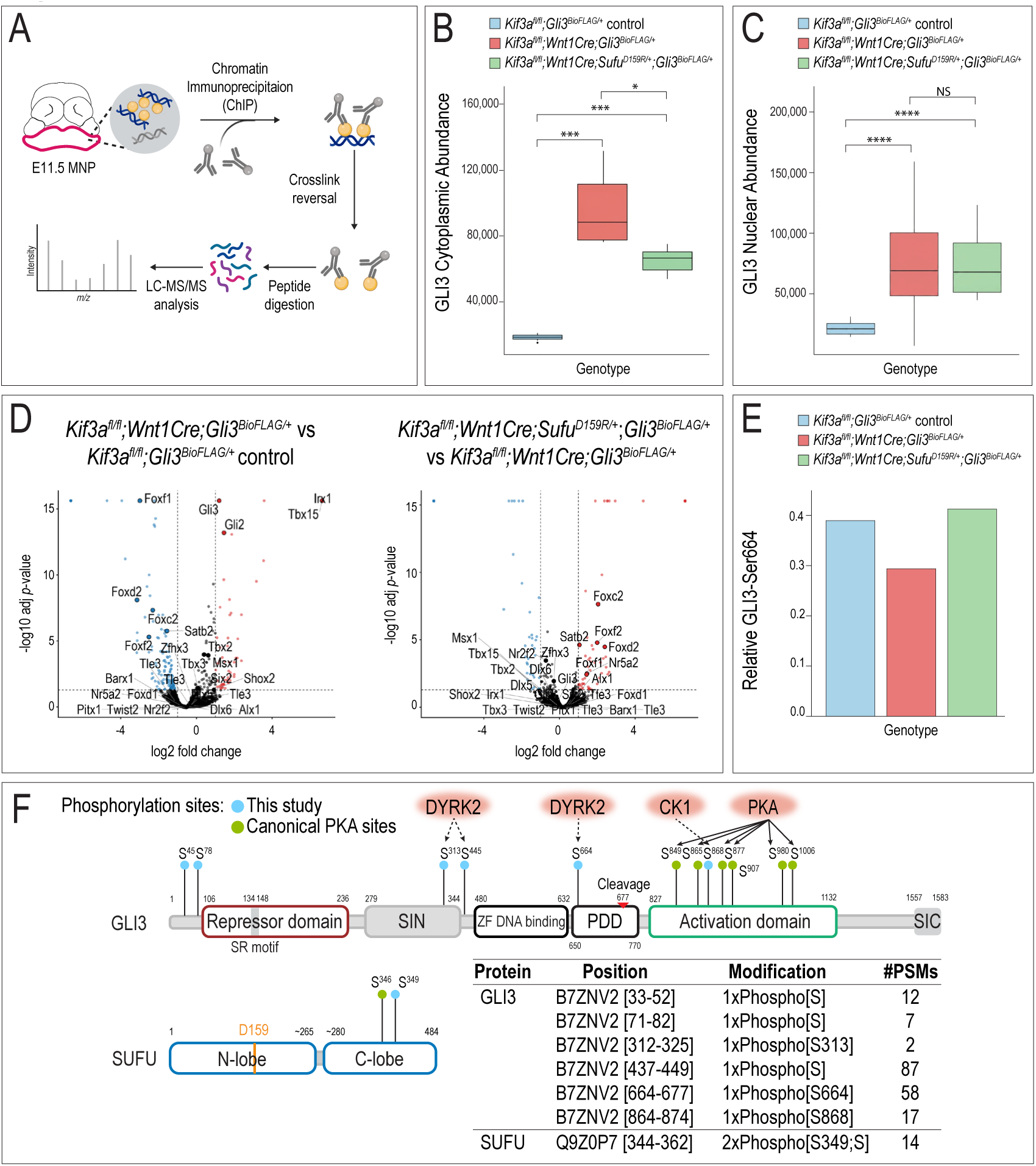
Introduction of *Sufu^D159R^* restores the GLI3-interactome and post-translational modifications (PTMs). (A) Schematic of the Rapid Immunoprecipitation Mass Spectrometry of Endogenous Proteins (RIME) workflow. (B, C) Quantification of GLI3 protein abundance in cytoplasmic and nuclear fractions. (D) Volcano plot showing differential protein abundances for GLI3 interactors highlighting statistically significant transcription factors (fold-change > 2, *adj p*-value < 0.05). (E) Ratio of phosphorylated to unphosphorylated GLI3 Ser664 measured using mass spectrometry. (F) Summary of phosphorylation events detected *in vivo* for GLI3 and SUFU. Table lists phosphorylation sites identified in this study (protein accession numbers used as reference), showing the peptide fragment position, the modification event and the number of peptide spectral matches (PSMs). Phosphorylation results were cross-referenced with UniProt to identify most likely modified residue when not determined by analysis pipeline. (\**p < 0.05*, \*\*\**p < 0.001*, \*\*\*\**p < 0.0001*, NS = not significant; PDD: processing

In addition to cofactor binding, post-translational modifications (PTMs) also alter GLI TF functionality (48, 49). GLI TFs are known to be phosphorylated, sumoylated, and acetylated (48). Phosphorylation clusters on GLI proteins are known to regulate both the formation of GLI repressors and their transcriptional activity (49). Analysis of proteomic data revealed phosphorylated peptide fragments and consistently recovered phosphorylation of GLI3-Ser664 (Fig. 7E), a site recently associated with cancer metastasis (50). Notably, the ratio of phosphorylated GLI3-Ser664 to total GLI3 from the peptide quantification data was reduced in *Kif3a^fl/fl^;Wnt1Cre;Gli3^BioFLAG/+^* compared to *Gli3^BioFLAG/+^* controls and increased in *Kif3a^fl/fl^;Wnt1Cre;Sufu^D159R/+^*;*Gli3^BioFLAG/+^* (Fig. 7E). In addition to RIME, we performed follow-up phospho-enrichment mass spectrometry on wild type tissue that confirmed Ser664 and detected additional phosphorylation events on GLI3 that conform to the consensus motifs of DYRK2 or CK1, suggesting potential regulation by these kinases (Fig. 7F, GLI3-blue circles). For SUFU, we identified a previously characterized protein kinase A (PKA) phosphorylation site Ser346 (51) (Fig. 7F, SUFU-green circles) and a novel site, Ser349 (Fig. 7F, SUFU-blue circles). These phospho-enrichment experiments failed to identify previously reported PKA target sites on GLI3 (52), presumably due to these modifications resulting in destabilization of GLI3 (Fig. 7F, GLI3-green circles). Together, these findings validated previous work which suggested that loss of cilia altered GLI3 PTMs (51, 53) and suggested that manipulating SUFU-GLI3 interactions exposed residues amenable to PTMs which contribute to modulating GLI3 transcriptional activity.

In summary, the phenotypic rescue of ciliopathic phenotypes associated with disrupting SUFU-GLI interactions, specifically via addition of the *Sufu^D159R^* allele, occurred via a mechanism that restored DNA binding and transcriptional regulatory activity of both GLI3A and GLI3R lost under ciliopathic conditions. These data further suggest that partially alleviating SUFU-GLI3 interactions bypassed the need for cilia and exposed residues on GLI3 that were essential for DNA binding, required for cofactor binding, and/or subject to PTM important for functionality. Importantly, unlike previous approaches to rescue ciliopathic phenotypes that exacerbated a subset of phenotypes (15, 16), introduction of the *Sufu^D159R^* allele partially rescued all facial phenotypes and restored transcriptional activity of both GLI3A and GLI3R. Together, these studies suggest targeting SUFU-GLI3 interactions as a potential therapeutic avenue for future treatment of ciliopathies.

## DISCUSSION

This study began with the premise that a deeper mechanistic understanding of the interaction between SUFU and GLI proteins might reveal a novel approach to rescue ciliopathic phenotypes. Using the *Kif3a^fl/fl^;Wnt1Cre* ciliopathic mouse model, our data indicate that interactions with SUFU were essential for transcriptional activation by GLI3A and repression by GLI3R during early facial development. Loss of primary cilia resulted in skewing of the SUFU-GLI3FL/SUFU-GLI3R ratio towards SUFU-GLI3FL, and loss of SUFU-GLI3 DNA binding at GLI3 target loci (Fig. 3B and Fig. 6). To avoid unintended phenotypic exacerbation previously observed when GLIA or GLIR were overexpressed (15, 16), we took advantage of the epistatic relationship between SUFU and the cilium (14, 32). Given the observed and previously reported upregulation of SUFU in ciliopathic mutants (15, 51), we introduced *Sufu^+/-^* or *Sufu^D159R^* alleles into a ciliopathic mouse model. While addition of either *Sufu^+/-^* or *Sufu^D159R/+^* alleles resulted in partially rescued midfacial phenotypes, the *Sufu^D159R/+^* allele was superior in restoration of mandibular phenotypes, both phenotypically and molecularly. Alteration of amino acid D159 to arginine restored the ability of SUFU-GLI complexes to bind to DNA and physically interact with transcriptional cofactors (Figs. 6-8). Together, these data suggest that targeting SUFU-GLI interactions is a plausible approach for the treatment of ciliopathies.

**Figure 8.**
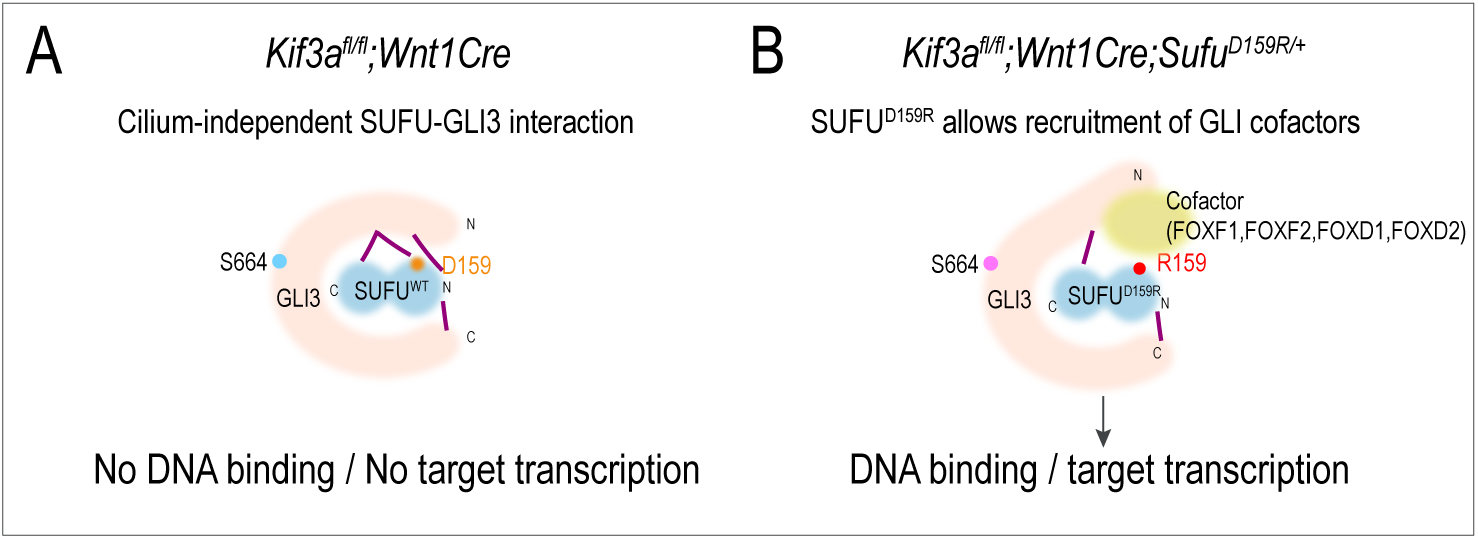
Proposed model for how SUFU^D159R^ restores GLI3 activity under ciliopathic conditions. Hypothesized model integrating our structural, biochemical, and transcriptomic data. In both *Kif3a^fl/fl^;Wnt1Cre* and *Kif3a^fl/fl^;Wnt1Cre;Sufu^D159R/+^* embryos SUFU remains bound to GLI3 through a cilium-independent interaction, with SUFU-GLI3 contacts indicated by purple lines. (A) In *Kif3a^fl/fl^;Wnt1Cre* ciliopathic mutants, wild-type SUFU interacts with GLI3 through the D159 (orange) contact within the GLI3 SIN domain; in this configuration GLI3 fails to bind DNA and does not activate transcriptional targets. (B) In *Kif3a^fl/fl^;Wnt1Cre;Sufu^D159R/+^* embryos, substitution of aspartic acid at position 159 with a basic arginine (R159, red) disrupts the D159/SIN contact and alters the conformation of the SUFU-GLI3 interaction. This altered conformation exposes GLI3 residues required for DNA binding and for recruitment of transcriptional cofactors, including FOX-family transcription factors (FOXF1, FOXF2, FOXD1, FOXD2; yellow), thereby restoring GLI3 DNA binding and target-gene transcription. A dynamically regulated GLI3 phosphorylation site, Ser664 (S664), is depicted on GLI3. Phosphorylation of S664 is reduced in the ciliopathic mutant (blue)and increased in the *Kif3a^fl/fl^;Wnt1Cre;Sufu^D159R/+^* (pink), illustrating that the SUFU-GLI3 interaction is further modulated by post-translational modification.

SUFU-GLI interactions are mediated by the SUFU interacting N-terminal domain (SIN), the SUFU interacting C-terminal domain (SIC), and the SUFU-binding repressor domain (SR), which is only present in GLI2 and GLI3. Interactions between SUFU and the SIN domain are required for GLI stabilization, cytoplasmic retention, ciliary-localization, and a target site for phosphorylation by multiple protein kinases (23, 25–28). Conversely interactions between the SIC domain and SUFU are required for its transcriptional activity (27). Under wild-type conditions, loss of interactions between SUFU and the GLI SIN domain, such as what occurs with SUFU^D159R^, results in GLI degradation (Fig. 3A). However, when SUFU^D159R^ was introduced on a ciliopathic background, the stability of, and interactions between GLI3 and SUFU were maintained, likely through interactions with the SIC and SR domains (Supplementary Fig. 2A). Our data support the conclusion that the SIC domain is critical and functionally important for GLI3 since introduction of the *Sufu^+/-^* allele failed to rescue MNP phenotypes to the same extent as *Sufu^D159R^*. We hypothesize that introduction of SUFU^D159R^ alleviated binding through the SIN domain, exposing key residues of the GLI3 protein required for the SUFU-GLI complex to bind to DNA, interact with other transcriptional cofactors, and/or restore expression of TFs such as the FOX TFs (Figs. 6,7,8). These results further suggest that under non-pathological conditions these events are cilia-dependent. More specifically, these data suggest that disrupting SUFU-GLI interactions under ciliopathic conditions exposes certain protein domains or residues that are post-translationally modified and/or required for interactions with requisite cofactors of GLI proteins (Figs. 7, 8). To test this hypothesis comprehensively, additional proteomic studies such as high-sensitivity mass spectrometry alongside mutagenesis strategies on isolated SUFU-GLI3 complexes and crystallography will be required.

SUFU has long been described as a negative regulator of the HH pathway, based largely on the gain-of-function phenotypes associated with *Sufu^-/-^* embryos (17, 41) and *in vitro* studies (18, 20). Moreover, SUFU recruits the SAP18-SIN3A corepressor complex to allow GLIR to repress transcriptional targets (54). Other studies, however, have suggested that SUFU is required for maximal HH activation (32), although the mechanisms of this role are not fully understood. Here, we show that SUFU interacts physically with both full-length GLI3A and truncated GLI3R isoforms *in vivo* to facilitate binding to DNA, indicating a role for SUFU in regulating the transcriptional activity of both GLI3A and GLI3R. Moreover, we observed SUFU preferentially bound to full-length GLI3FL (Fig. 1B), and SUFU-mediated GLI3A activity was affected more than SUFU-mediated GLI3R activity in the absence of primary cilia (Fig. 5B), supporting the hypothesis that SUFU activity is essential for both activation and repression of GLI targets.

Several ciliopathic phenotypes result from impaired GLI activity (15, 16). In this study, we not only identified novel targets downstream of GLI3 (Table S6), but also found that their expression was lost under ciliopathic conditions, and subsequently restored with the introduction of SUFU^D159R^ (Fig. 5). Motif analysis (Fig. 6D-G) as well as RIME (Fig. 7) suggest that some of these targets may function as GLI cofactors (e.g., FOX, Homeobox, and T-box factors) that play essential roles in craniofacial development, specifically members of the FOX family (FOXC, FOXD, FOXF) (55–58). Notably, RIME analysis revealed that the partially restored phenotypes observed in our study correlated with restored physical interactions between GLI3 and FOX family TFs (Fig. 7D). Future studies will focus on delineating the specific cofactors that interact with GLI3A versus GLI3R to further understand their distinct regulatory mechanisms.

In addition to interactions with key cofactors, GLI activity also relies on post-translational modifications. Phosphorylation at sites targeted by PKA (11, 13, 52) and further phosphorylation by CK1/GSK3β at adjacent sites (59) induces GLI processing to the repressor, whereas phosphorylation at sites N-terminal to zinc-finger domain are responsible for GLI activation (52). A recent study also showed that phosphorylation by DYRK2 positively regulates GLI2 and GLI3 (60) (Fig. 7F). RIME analysis herein suggests that introduction of SUFU^D159R^ alters the conformation of the SUFU-GLI complex in such a way that exposes key residues for post-translational modification. For example, phosphorylation of GLI3-Ser664 was reduced in *Kif3a^fl/fl^;Wnt1Cre* and restored in *Kif3a^fl/fl^;Wnt1Cre;Sufu^D159R/+^*, raising the possibility that residues requiring post-translational modification for GLI function are masked or altered by aberrant SUFU interactions in the absence of cilia. Therefore, tuning kinases or phosphatases responsible for these sites pharmacologically could potentially mimic the effect of modulating the SUFU-GLI3 interface. However, future work using high-resolution phosphoproteomics or structural studies will be needed to define the full extent of PTM regulation and how it integrates with cofactor recruitment and DNA binding.

Finally, the *Sufu^D159R^* allele was generated on a C57BL/6J background, whereas the *Wnt1Cre* and *Kif3a^fl/fl^* lines are maintained on a mixed background; the triple-transgenic crosses therefore introduce both the *Sufu^D159R^* allele and a shift toward the C57BL/6J background. We cannot fully exclude the possibility that background modifiers contribute to the partial rescue. However, the molecular data (CUT&RUN, RNA-seq, and RIME) support specific restoration of GLI3 DNA binding, target-gene expression, and cofactor interactions that are mechanistically consistent with disruption of the SUFU-GLI3 interface rather than with background effects. Moreover, the *Sufu*-null allele, which is on a C57BL/6J background and therefore introduces a comparable background shift, does not rescue mandibular phenotypes to the same degree as *Sufu^D159R^*, arguing against a purely background-driven effect.

Due to a limited understanding of ciliary function and the pleiotropic nature of ciliopathies, treatment strategies for these disorders are extremely limited. Currently the only approach focuses on symptomatic relief, rather than addressing the root cause of these disorders. In the case of craniofacial skeletal phenotypes in human patients, current surgical treatments, such as distraction osteogenesis for micrognathia, are highly invasive and have difficult recoveries. Any marginal improvement of phenotypes that could negate the need for these painful surgeries would be highly significant for patients. The data presented herein not only shed light on the role for cilia in GLI activity but also suggests that a better understanding of the structural interactions between SUFU and GLI, and their concomitant post-translational modifications (61, 62), could guide the development of pharmaceutical strategies aimed at curbing the aberrant activities of GLI in ciliopathies. Currently, however, these findings represent a genetic proof of concept rather than a therapeutic strategy. Complete disruption of the SUFU-GLI interface is deleterious. While homozygous *Sufu^D159R/D159R^* embryos are embryonic lethal, germline loss of SUFU in humans predisposes to medulloblastoma and other HH-driven tumors (23, 63, 64). Because SUFU regulates GLI activity across many tissues, systemic inhibition of SUFU would be expected to have on-target teratogenic and tumorigenic effects beyond the craniofacial structures examined here. The partial rescue achieved with a single *Sufu^D159R^* allele indicates that only a modest, tunable reduction in SUFU-GLI interaction is required. Any translational application would demand spatially and temporally restricted, dose-controlled modulation of the interface. Thus, further investigation will be required before these findings can be considered for clinical application.

## MATERIALS AND METHODS

### Structural modeling of the SUFU-GLI3 complex

Complexes of mouse SUFU with full-length GLI3 were predicted using AlphaFold3 (33) on the AlphaFold Server. Two jobs were submitted: full-length mouse SUFU (UniProt Q9Z0P7; 484 aa) with full-length mouse GLI3 (UniProt Q61602; 1,583 aa), and an otherwise identical job in which SUFU Asp159 was substituted with Arg (D159R), corresponding to the *Sufu^D159R^* allele. Neither chain was truncated, so the intrinsically disordered regions flanking the GLI3 zinc-finger DNA-binding domain were retained. Five Zn^2+^ ions were included as ligands to support folding of the five-zinc-finger DNA-binding domain, and structural templates and automatic multiple sequence alignment generation were enabled. Each job used a single seed (seed 5) and produced five models; the top-ranked model was used for all analyses and the remaining models were checked for consistency. Interface quality was assessed from ipTM, the SUFU-GLI3 chain-pair ipTM, the predicted aligned error (PAE), and per-residue pLDDT. Global pTM was not used for interpretation because full-length GLI3 is largely intrinsically disordered. Inter-chain contacts were taken directly from the AlphaFold3 token-level contact-probability matrix rather than from a distance cutoff. All SUFU-GLI3 token pairs with a contact probability ≥ 0.5 were retained together with their symmetrized PAE, and per-residue pLDDT was read from the Cα records of the mmCIF model. Contacts were plotted as Circos-style chord diagrams using circlize in R. Each chord links one SUFU residue to one GLI3 residue and is colored by contact probability. Identical parameters were applied to both models, so differences in chord density reflect differences in the predicted interface.

### Transgenic mice

*Sufu^D159R^* mice were generated by CRISPR targeting in the Transgenic Animal and Genome Editing (TAGE) facility at CCHMC and maintained on a C57BL/6J background. *Wnt1Cre* (JAX, 003829) and *Sufu-null* (JAX, 007859) mice were obtained from Jackson Laboratory. *Kif3a^fl/fl^* mice were obtained from Dr. Bradley Yoder, University of Alabama at Birmingham. *Gli3^BioFLAG^* knockin mice were obtained from Dr. Kevin Peterson (JAX, 026135). *Gli3^BioFLAG^* mice harbor an N-terminal 3xFLAG epitope for use with anti-FLAG antibodies. Mice with *Gli3^BioFLAG^* allele were used for GLI3 CUT&RUN and immunoprecipitation in this study. All the mice were maintained by Veterinary Services of Cincinnati Children’s Hospital Medical Center with IACUC approval. Except *Sufu^D159R^* mice, all other double or triple transgenic lines were maintained on a mixed background.

### Imaging/Histology

E9.5 or E15.5 embryos were harvested and fixed in Bouin’s fixative overnight for whole mount imaging. Images were taken with Leica M165FC microscope and Leica LAS X software. For Safranin-O staining, tissue was processed for paraffin embedding and staining procedures were followed as described (http://www.ihcworld.com/_protocols/special_stains/safranin_o.htm). All histological and imaging analyses were performed on a minimum of n = 3 embryos per genotype.

### SpDamID-seq

O9-1 and MK4 cells stably expressing the TetOn transactivator (pLenti CMV rtTA3 Blast, Addgene 26429) were seeded on 6-well plates at approximately 80% confluence and transfected with Lipofectamine 3000 (Invitrogen) and 1.0 μg/well of the pBI-Tet or pCMV-3X-FLAG vectors containing the indicated Split DamID fusion protein pairs. Plasmids used for transient transfections were produced in competent Dam^-/-^; Dcm^-/-^ bacteria (NEB C2925H) to avoid amplification of methylated plasmid DNA. Twenty-four hours after transfection, medium was changed and 1 μM doxycycline and 2 μM 4-hydroxytamoxifen (4-OHT) were added to induce expression and nuclear translocation of the ERT2-containing fusion proteins. Forty-eight hours post-transfection the DNA was extracted using Purelink Genomic DNA mini kit (Invitrogen K182001). One μg of genomic DNA was digested overnight with DpnI, adaptors ligated onto the digested DNA ends and the library amplified using primers as in (37). Split DamID-seq reads were aligned to the GRCm38/mm10 with Bowtie2 and processed with the damidseq_pipeline (65), which computes GATC-fragment-resolution, RPM-normalized log2 ratios of each Dam-fusion sample over its matched unfused Dam-only control using a genome-wide GATC fragment reference. Each cell line (O9-1 and MK4), GLI3 isoform (GLI3FL-AM or GLI3R-AM co-expressed with D-SUFU), and induction condition (± doxycycline) was processed independently against its matched Dam-only control. Methylation-enriched peaks were called with MACS2 using the Dam-only sample as the control; high-confidence peaks were defined as MACS2 peaks overlapping a genomic GATC fragment with a false-discovery rate of *q* < 0.01. Peaks were annotated to genes with ChIPseeker. The induced (+Dox) peak sets were used for all reported analyses. Gene ontology (GO) was analyzed by ToppGen Suite (https://toppgene.cchmc.org).

### Bulk RNA-sequencing and analysis

For the E9.5 dataset, RNA was isolated from whole heads of wild-type, *Sufu^-/-^*, and *Sufu^D159R/D159R^* embryos (≥3 biological replicates per genotype). For the E11.5 dataset, RNA was isolated from frontonasal (FNP) and mandibular (MNP) prominences of wild-type, *Kif3a^fl/fl^;Wnt1Cre*, *Kif3a^fl/fl^;Wnt1Cre;Sufu^+/-^*, and *Kif3a^fl/fl^;Wnt1Cre;Sufu^D159R/+^* embryos (n = 3 per genotype per prominence). Total RNA was isolated with the RNAqueous-Micro Total RNA Isolation Kit (Thermo Fisher, AM1931). Poly-A-selected libraries were prepared with the Illumina TruSeq stranded mRNA kit and sequenced on an Illumina NovaSeq 6000, yielding ≥ 20 million 100-bp paired-end reads per sample.

For the E9.5 dataset, reads were processed with the nf-core/RNAseq pipeline and quantified to gene level with salmon against the Ensembl GRCm39/mm39 reference; because samples spanned multiple sequencing batches, counts were batch-corrected with ComBat-Seq, and both differential expression and principal-component analysis (Fig. 2F) were performed on the corrected counts. For the E11.5 dataset, reads were trimmed with fastp, aligned to the GRCm39/mm39 reference with STAR, and counted with featureCounts (paired-end, read-pair counting, automatic strand detection).

Differential expression was performed with DESeq2 using genotype as the sole factor. For the E9.5 dataset, genes with a Benjamini–Hochberg adjusted p < 0.05 and the absolute value of log2 fold-change ≥ 1 were called differentially expressed (1,655 genes for *Sufu^-/-^* versus wild-type; 2,264 for *Sufu^D159R/D159R^* versus wild-type). For the E11.5 dataset, expression was analyzed separately per prominence using a pairwise-subset strategy, and log2 fold-changes for genes with adjusted p < 0.05 were consider significant. A gene was defined as dysregulated in the ciliopathic mutant if it had adjusted p < 0.05 and the absolute value of log2 fold-change > 1 in the *Kif3a^fl/fl^;Wnt1Cre* versus wild-type (22 genes in the FNP, 39 in the MNP).

For the E11.5 rescue analysis, a dysregulated gene was called restored by a rescue allele if that allele produced a significant change relative to *Kif3a^fl/fl^;Wnt1Cre* (adjusted p < 0.05 in rescue versus mutant) in the direction opposite to the mutant effect. For each dysregulated gene, the degree of restoration (the fraction of the mutant dysregulation corrected) was defined as 1 − (|log2 fold-change of rescue versus wild-type| ÷ |log2 fold-change of mutant versus wild-type|), where 1 indicates full return to wild-type, 0 no correction, and negative values displacement further from wild-type than the mutant. The two rescue alleles, paired by gene, were compared with a two-sided exact Wilcoxon signed-rank test. For heatmaps, DESeq2 size-factor normalized counts were variance-stabilized (vst, blind = FALSE) and z-scored per gene across the twelve samples of each prominence; genes were clustered by Euclidean distance (complete linkage) and columns ordered by genotype without clustering. Only mutant-dysregulated genes restored by at least one allele were shown; one unannotated model gene (LOC118568416) was excluded from the MNP panel, giving final sets of 10 (FNP) and 14 (MNP) genes.

### CUT&RUN-seq

CUT&RUN-seq was performed based on the manufacturer’s protocol (EpiCypher CUTANA CUT&RUN Protocol v1.7). Briefly, E11.5 mouse MNPs (control *Kif3a^fl/fl^;Gli3^BioFLAG/^*: n = 3; *Kif3a^fl/fl^;Wnt1Cre;Gli3^BioFLAG/+^*: n = 2; *Kif3a^fl/fl^;Wnt1Cre;Sufu^D159R/+^;Gli3^BioFLAG/+^*: n = 3) were harvested in ice cold PBS and digested with 100 μg/mL Liberase (Sigma, 5401119001) in PBS at 37°C for 5 minutes. 250,000-400,000 cells were then incubated with activated ConA beads (EpiCypher, 21-1411) at 4°C for 30 minutes, followed by antibody incubation (FLAG-M2 1:100 (Sigma, F3165), SUFU 1:100 (Proteintech, 26759-1-AP)), diluted in 0.025% Digitonin buffer containing 2 mM EDTA on a nutator at 4°C overnight. Samples were washed three times in Digitonin buffer by gentle pipetting, incubated with secondary antibodies for 30 minutes at 4 °C, washed, incubated with pAG-MNase (EpiCypher, 15-1016) for 30 minutes at 4 °C, followed by three washes. The MNase reaction was activated by 2 mM CaCl_2_ in Digitonin buffer at 0 °C ice bath for 1 hour and terminated by stop buffer. DNA fragments were purified by phenol-chloroform extraction and libraries were generated using NEBNext Ultra II DNA Library Prep Kit (NEB, E7645S). AMPure XP beads (Beckman Coulter, A63881) were used for adapter removal and the final clean-up for libraries. Samples were sequenced on an Illumina NextSeq 500 instrument using PE-75 bp to a depth of 3-5 million reads. CUT&RUN sequencing data were analyzed on the Galaxy platform (usegalaxy.org) (66). Paired-end reads were aligned to the mouse genome (GRCm38/mm10) with Bowtie2 using default parameters and the built-in mm10 index. Peaks were called with MACS2 in paired-end mode (BAMPE), pooling replicate (treatment) libraries and comparing them against pooled control (IgG) libraries. A peak model was built (mfold bounds 5–50; bandwidth 300 bp), peaks were detected by q-value at a FDR cutoff of q < 0.05 in narrow-peak mode. Peaks were annotated by HOMER.

### Western blot, transfection, immunoprecipitation, and immunostaining

For western blot (WB) analysis of early-stage embryos, protein lysate was prepared from E9.5 whole embryos or E11.5 facial prominences in RIPA buffer with protease and phosphatase inhibitors (Thermo Fisher Scientific, 78440). For immunoprecipitation (IP), MK4 cells (mouse embryonic kidney cells, Dr. Steve Potter, CCHMC) were transfected at ∼70% confluency using Lipofectamine 3000 (Invitrogen, L3000), with different combinations of Gli and Sufu constructs [mouse full-length Gli2, full-length Gli3, or truncated Gli3 repressor (1–699) that were cloned into p3xFLAG-CMV-7.1; EGFP-tagged mouse Sufu (Addgene, 65431); EGFP-tagged Sufu^D159R^ that was generated by Q5® Site-Directed Mutagenesis Kit (NEB, E0554S)]. 24 hours after transfection, cells were lysed in RIPA buffer with protease and phosphatase inhibitors and ∼500 μg protein was used for IP. A subset of the transfected cells was fixed in 4% PFA for immunofluorescent (IF) staining. The following primary and secondary antibodies were used for WB, IP or IF: anti-GLI3 (1:1000; R&D Systems, AF3690), anti-SUFU (1:1000; Proteintech, 26759-1-AP), anti-FLAG-M2 (1:1000 for WB, 1:500 for IF, 5 μg for IP; Sigma, F3165), anti-VINCULIN (1:2000; Santa Cruz Biotechnology, sc-73614), IRDye® 800CW Donkey anti-Rabbit IgG (1:2000; LI-COR 926-32213), IRDye® 680RD Donkey anti-Mouse IgG (1:2000; LI-COR 925-68072). Images were taken by LI-COR Odyssy® DLx. Densitometry was done by ImageJ. All western blot and co-immunoprecipitation experiments were performed with a minimum of n = 2 independent biological replicates.

### RNAscope *In Situ* Hybridization Assay and quantification

Embryos (E10.5 wild-type, *Kif3a^fl/fl^;Wnt1Cre* and *Kif3a^fl/fl^;Wnt1Cre;Sufu^D159R/+^*: n = 3) were fixed in 4% paraformaldehyde diluted in phosphate buffer at room temperature for 2 hours. Embryos were dehydrated through an ethanol series, washed in xylene, embedded in paraffin, and sectioned at 6 µm thickness. *Foxf2* (Advanced Cell Diagnostics 473391) transcripts were detected using the RNAscope Multiplex Fluorescent V2 kit (ACD, Newark, CA, USA) per manufacturer’s instructions. Hydrogen peroxide treatment was reduced to 5 minutes. Signal development for each probe was done using fluorescein (Akoya Biosciences NEL74100KT, Marlborough, MA, USA) or Cyanine 3 (Akoya Biosciences NEL744001KT, Marlborough, MA, USA) diluted 1:500 in RNAscope Multiplex TSA buffer. Slides were counterstained and mounted with Prolong Gold with DAPI (Invitrogen, P36931) and imaged using a Leica DM5000B upright microscope system.

### Mass spectrometry and analysis

To identify GLI3-interacting proteins, we performed a modified Rapid Immunoprecipitation Mass Spectrometry of Endogenous (RIME) protocol (47) on E11.5 MNPs (*Kif3a^fl/fl^;Gli3^BioFLAG/+^* control, n = 3; *Kif3a^fl/fl^;Wnt1Cre;Sufu^D159R/+^;Gli3^BioFLAG/+^*, n = 12; *Kif3a^fl/fl^;Wnt1Cre; Gli3^BioFLAG/+^*, n = 15) that were snap frozen in liquid nitrogen and stored at -80 °C until use. Briefly, intact samples were crosslinked with 1% formaldehyde for 8 minutes at room temperature, quenched with glycine (0.1M), and washed twice with ice cold 1X phosphate buffered saline, followed by nuclear-cytoplasmic fractionation. The cytoplasmic fraction was snap frozen and stored at -80 °C. The nuclear fraction was further processed for sonication and immunoprecipitated overnight with mouse monoclonal anti-FLAG antibody (Sigma-Aldrich F3165) conjugated to sheep anti-mouse IgG magnetic beads (Thermo Fisher 11201D). Both IgG controls and mock anti-FLAG controls were run in parallel on lysates prepared from wild-type C57BL/6J MNPs. Following the immunoprecipitation steps in the RIME protocol, samples were on-bead digested and peptides were desalted by solid-phase extraction using C18 ultra micro spin columns. Peptides were dried with a vacuum centrifuge and were stored at -80°C. Samples were then run using data independent acquisition (DIA) on a Thermo Orbitrap Astral coupled to a Vanquish Neo liquid chromatography system using an EASY-spray column (PepMap Neo C18, 75 μm x 150 mm, #ES75150PN) in the Mass Spectrometry and Protein Chemistry Service at The Jackson Laboratory. The 25-minute gradient using a flow rate of 0.6 μL/min and all mass spectrometry method settings were as described previously (47, 67).

The RAW files from the Thermo Orbitrap Astral underwent automatic processing on the Thermo Ardia Server and were analyzed using Proteome Discoverer 3.1 with CHIMERYS on the associated workstation. This analysis was performed as described previously (67) with the addition of a dynamic phosphorylation of serine, threonine, and tyrosine to those search settings. First, each genotype (*Kif3a^fl/fl^;Gli3^BioFLAG/+^*, *Kif3a^fl/fl^;Wnt1Cre;Gli3^BioFLAG/+^*, and *Kif3a^fl/fl^;Wnt1Cre;Sufu^D159R/+^;Gli3^BioFLAG/+^*) was filtered against IgG and mock controls by applying the following cut-offs: False Discovery Rate < 0.05, peptide spectral matches ≥ 6, and Abundance Ratio (log2) ≥ 2 versus both IgG and mock controls. A final set of 6817 non-redundant GLI3 enriched proteins was determined by filtering across 3 groups with the following cut-offs: FDR < 0.05, Peptide Spectral Matches ≥ 6, and Abundance Ratio (log2) ≥ 1. Differential analysis was performed between groups and significantly enriched proteins were determined by applying an additional adjusted p-value cut-off < 0.05. Additionally, 1112 phospho-peptides were identified within the RIME dataset by filtering for phosphorylation modification and Abundance Ratio (log2) ≥ 1 in at least one of the groups versus both mock and IgG controls.

In addition to the immunoprecipitation samples, peptide samples from protein lysates were prepared from *Kif3a^fl/fl^;Gli3^BioFLAG/+^* E11.5 heads as described previously with the addition of 1X PhosSTOP (Roche, #4906845001), followed by PTMScan IMAC Phospho-enrichment according to manufacturer protocol (Cell Signaling Technologies, #20432). These peptides were then desalted by solid-phase extraction using C18 ultra micro spin columns as above and were dried with a vacuum centrifuge.

## DATA AVAILABILITY

Sequencing data have been deposited in GEO repository under accession codes GSE301704 and GSE301705 for CUT&RUN, GSE301706, GSE301707 and GSE348431 for RNA-seq, and GSE301638 for Split DamID-seq.

## ACKNOWLEDGEMENTS

We would like to thank the Mass Spectrometry and Protein Chemistry Service within Protein Sciences at The Jackson Laboratory for their support with the mass spectrometry analysis performed in this study. The mass spectrometry method development in this project was also supported by mass spectrometry instrumentation from the National Institutes of Health S10-OD026816 award. This work was supported by the National Institutes of Health and the National Institute of Dental and Craniofacial Research (R35 DE027557 to S.A.B. and R01 DE031750 to K.A.P. and S.A.B.)

## Supplementary Tables

**Table S1.** AlphaFold3 confidence metrics and inter-chain contacts for GLI3 with SUFU^WT^ and SUFU^D159R^.

**Table S2.** Gene annotations of peaks from Split DamID.

**Table S3.** Differentially expressed genes between *Sufu^-/-^* or *Sufu^D159R/D159R^* and wild-type embryos.

**Table S4.** Differentially expressed genes common in *Sufu^-/-^* and *Sufu^D159R/D159R^* embryos, and their overlap with GLI targets.

**Table S5**. List of differentially expressed genes in *Kif3a^fl/fl^;Wnt1Cre* compared to wild-type and *Kif3a^fl/fl^;Wnt1Cre*;*Sufu^+/-^* or *Kif3a^fl/fl^;Wnt1Cre*;*Sufu^D159R/+^* compared to *Kif3a^fl/fl^;Wnt1Cre*.

**Table S6**. Gene annotations of peaks from CUT&RUN and novel GLI3 targets identified in this study.

**Table S7**. RIME analysis identifying GLI3 associated proteins and phosphorylated residues.

